# The impact of SNP density on quantitative genetic analyses of body size traits in a wild population of Soay sheep

**DOI:** 10.1101/2022.03.07.483376

**Authors:** Caelinn James, Josephine M. Pemberton, Pau Navarro, Sara Knott

**Author notes:** Corresponding author Corresponding author mailing address: Caelinn James, Ashworth Laboratories, Kings Buildings, Charlotte Auerbach Road, Edinburgh EH9 3FL Corresponding author.

## Abstract

Understanding the genetic architecture underpinning quantitative traits in wild populations is pivotal to understanding the processes behind trait evolution. The ‘animal model’ is a popular method for estimating quantitative genetic parameters such as heritability and genetic correlation and involves fitting an estimate of relatedness between individuals in the study population. Genotypes at genome-wide markers can be used to estimate relatedness; however, relatedness estimates vary with marker density, potentially affecting results. Increasing density of markers is also expected to increase the power to detect quantitative trait loci (QTL). In order to understand how the density of genetic markers affects the results of quantitative genetic analyses, we estimated heritability and performed genome-wide association studies (GWAS) on five body size traits in an unmanaged population of Soay sheep using two different SNP densities: a dataset of 37,037 genotyped SNPs, and an imputed dataset of 417,373 SNPs. Heritability estimates did not differ between the two SNP densities, but the high-density imputed SNP dataset revealed four new SNP-trait associations that were not found with the lower density dataset, as well as confirming all previously-found QTL. We also demonstrated that fitting fixed and random effects in the same step as performing GWAS is a more powerful approach than pre-correcting for covariates in a separate model.

## Introduction

Investigating the genetic architecture behind heritable traits is key to understanding the biological diversity of wild populations. If we know the number of loci influencing a trait and their effect size, we can better understand the evolutionary processes that underpin traits, improve inferences about trait evolution, and understand micro-evolutionary dynamics that occur due to environmental change (Barton and Keightley 2002). Most quantitative genetic research in animals is carried out in artificial populations; either domestic, agricultural or laboratory. Such populations experience controlled environmental conditions which make it easier to account for environmental factors when studying the effect of genetic variants on phenotypic variation. However, given that environmental factors can influence the phenotype of a quantitative trait (Charmantier et al. 2014), and that the presence of genotype-by-environment interactions can cause additive genetic variance to differ between environmental conditions, studies on artificial populations arguably cannot be fully extrapolated to wild populations (Kruuk et al. 2008). Therefore, it is important to also study quantitative traits in wild populations in their natural habitats. There is a wealth of quantitative genetics research in human populations (for examples, see Manolio et al. 2009; Kang et al. 2010; Yang et al. 2010; Zaitlen et al. 2013; Locke et al. 2015; Xia et al. 2016; Xia et al. 2021), but humans also arguably experience a more buffered environment than wild populations and inferences in wild populations are lacking in comparison.

When using molecular markers to inform quantitative genetic analyses, the results are dependent on the genetic polymorphisms used in the analysis: low numbers of polymorphisms can result in downwardly biased heritability estimates and regions containing causal variants may not appear as significant if there are no genotyped polymorphisms in linkage disequilibrium (LD) with the causal variant. Increasing the density of genotyped polymorphisms means they are more likely to be in LD with causal variants for the trait of interest, either by being physically closer to the causal variants or by matching the allele frequency of the causal variants more accurately. However, increasing the number of genotyped markers means larger, denser genotyping arrays with costs increasing with density. For commonly studied species, high-density arrays are more affordable due to high demand, but for more niche species, including wild populations, large genotyping arrays are often unaffordable. Genotyping-by-sequencing, e.g. ddRAD (Peterson et al. 2012) is a potentially useful alternative for upscaling SNP density, though the combination of bioinformatics and samples sizes required in quantitative genetic research means that this approach is not yet in widespread use.

As an alternative to expensive high-density genotyping, genotype imputation can be used to increase the number of variants analysed (Burdick et al. 2006). Imputation involves predicting genotypes at untyped SNPs in a ‘target’ population using a subset of the study population – or more generally a “reference” population – genotyped at a higher density, either through a high density SNP array or by genotyping-by sequencing. The genotypes at these untyped SNPs for individuals in the target population are inferred using their genotypes at typed markers and taking advantage of existing linkage disequilibrium (LD) between SNPs. Pedigree information can also be used to increase the accuracy of the imputation by identifying haplotype blocks that are identical by descent (Burdick et al. 2006).

The Soay sheep (*Ovis aries*) of St Kilda are a primitive, unmanaged breed of sheep that have been the focus of a longitudinal, individual-based study since 1985 (Clutton-Brock and Pemberton 2003). As part of the study, morphometric, life history and environmental data are collected, DNA samples are collected, and a pedigree has been constructed using observation and genetic parentage inference. 7630 sheep have been genotyped on the Ovine SNP50 Illumina Beadchip, on which 37,037 SNPs are autosomal and polymorphic in this population. In addition, 188 individuals have been genotyped on the Ovine Infinum HD Beadchip, which contains 419,281 autosomal SNPs that are polymorphic in the population – this has allowed for imputation of the remaining sheep to this higher density (Stoffel et al. 2021).

In this study we performed a direct comparison of heritability estimates and GWAS associations between the lower density SNP data and the imputed high density SNP data in the Soay population, focusing on five body size traits in neonates, lambs and adults. We performed GWAS by fitting fixed and random effects in the same step as testing for SNP-trait associations, which has the advantage of correctly propagating error throughout the analysis, reducing the chance of false positive results and increasing power by disentangling potential correlations. We also carried out a two-step GWAS approach previously used on a smaller sample size of the Soays (Bérénos et al. 2015) to investigate whether any SNP-trait associations identified using our approach were due to the increased sample size or due to the different methodology (single-step vs. two-step GWAS).

Our aims were as follows:

1. To determine whether the increased density of SNPs changes the heritability estimates of the traits.
2. To determine whether the imputed SNP data enables the identification of new SNP-trait associations via GWAS.
3. To compare a single-step GWAS methodology with the two-step approach previously used on the study population.

Whilst we have used the Soay sheep as our study population, we believe that our objectives are also relevant for other wild populations. Keeping costs down is important for all research groups, and we aim to show that imputation is a way to do so whilst improving the power results of quantitative genetic analyses. We also intend to highlight the benefit of fitting fixed and random effects whilst performing GWAS instead of pre-correcting.

## METHODS

### Phenotypic data

The sheep are ear-tagged when they are first captured which allows for reidentification for life. We focused on five body size traits in three age groups: neonates, lambs, and adults. Of the five traits, three (weight, foreleg length and hindleg length) are live measures, recorded in April for neonates and in August for lambs and adults. The remaining two traits (metacarpal length and jaw length) are post mortem measures taken from skeletal material. Both birth and August weight are measured to the nearest 0.1kg, whilst the remaining traits are all measured to the nearest mm. A detailed description of trait measurements can be found in Beraldi et al. (2007).

We defined neonates as individuals who were caught and weighed between two and ten days after birth – birth weight was the only trait recorded for this age group. Lambs were classed as individuals who had phenotypic data recorded in the August of their birth year for the live traits, and as individuals who died before 14 months of age for the post mortem measures. Individuals were classed as adults if they had August phenotypic data recorded at least two years after birth, or if they died after 26 months of age for post mortem measures. We chose not to analyse yearling data due to the small sample sizes in comparison to the other age classes, which is due to high first winter mortality.

### Genetic data

Most of the sheep in our study population have been genotyped using the Ovine SNP50 Illumina BeadChip, which targets 54,241 SNPs across the sheep genome. After removing SNPs which failed quality control standards (minor allele frequency (MAF) > 0.001, call rate > 0.99, deviation from Hardy-Weinberg Equilibrium P > 1e-05) and individuals with a call rate < 0.95, 39,368 polymorphic variants remained for 7630 individuals (3643 female, 3987 male). See Bérénos et al. (2014) for information on genetic sampling protocol and marker characteristics).

Of these 7630 individuals, 188 have also been genotyped using the Ovine Infinium HD SNP BeadChip which targets 606,066 SNPs. This has allowed for the low density genotypes to be imputed to the higher density using AlphaImpute, which combines shared haplotype and pedigree information for phasing and genotype imputation (Hickey et al. 2012) (see Stoffel et al. (2021) for information on imputation and quality control). We used imputed genotype “hard” calls (rather than genotype probabilities) in downstream analyses. After filtering SNPs that failed quality control standards, 419,281 autosomal SNPs remained for 7621 individuals (3639 females, 3982 males).

Both the 50K SNP data and the imputed SNP data are mapped to the OAR_v3.1 genome assembly.

### Narrow sense heritability estimation

We used animal models to partition the phenotypic variance for each trait in each age class into genetic and non-genetic variance components. Fixed and random effects were fitted for all models, with the effects differing between traits and age classes (Table 1). We implemented these analyses in DISSECT (Canela-Xandri et al. 2015) using the following model:

**Table 1.** Number of individuals and records, fixed and random effects fitted in each trait x age class model in addition to the GRM. The same individuals and records were used for both heritability estimates and for GWAS.

| Age | Trait | No. individuals | No. records | Fixed effects | Random effects |
| --- | --- | --- | --- | --- | --- |
| Neonate | Birth weight | 2678 | 2678 | Sex | Year of birth |
|  |  |  |  | Litter size | Mother ID |
|  |  |  |  | Population size year before birth |  |
|  |  |  |  | Age of mother (quadratic) |  |
|  |  |  |  | Ordinal date of birth |  |
|  |  |  |  | Age (days) |  |
| Lamb | Weight | 2228 | 2228 | Sex | Year of birth |
|  |  |  |  | Litter size | Mother ID |
|  |  |  |  | Population size | Permanent environment |
|  |  |  |  | Age (days) |  |
|  | Foreleg | 2284 | 2284 | Sex | Year of birth |
|  |  |  |  | Litter size | Mother ID |
|  |  |  |  | Population size | Permanent environment |
|  |  |  |  | Age (days) |  |
|  | Hindleg | 2349 | 2349 | Sex | Year of birth |
|  |  |  |  | Litter size | Mother ID |
|  |  |  |  | Population size | Permanent environment |
|  |  |  |  | Age (days) |  |
|  | Metacarpal | 2059 | 2059 | Sex | Year of birth |
|  |  |  |  | Litter size | Mother ID |
|  |  |  |  | Age at death (months) |  |
|  | Jaw | 2113 | 2113 | Sex | Year of birth |
|  |  |  |  | Litter size | Mother ID |
|  |  |  |  | Age at death (months) |  |
| Adult | Weight | 1152 | 3553 | Sex | Year of capture |
|  |  |  |  | Population size | Permanent environment |
|  |  |  |  | Age (years) |  |
|  | Foreleg | 1121 | 3331 | Sex | Year of capture |
|  |  |  |  | Population size | Permanent environment |
|  |  |  |  | Age (years) |  |
|  | Hindleg | 1135 | 3444 | Sex | Year of capture |
|  |  |  |  | Population size | Permanent environment |
|  |  |  |  | Age (years) |  |
|  | Metacarpal | 945 | 945 | Sex | Year of birth |
|  |  |  |  | Age at death (years) |  |
|  | Jaw | 991 | 991 | Sex | Year of birth |
|  |  |  |  | Age at death (years) |  |

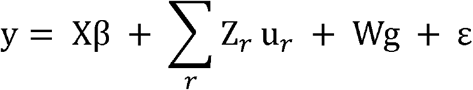

where y is the vector of phenotypic values; X is a design matrix linking individual records with the vector of fixed effects β, Z_r_ is an incidence matrix that relates the random effect r to the individual records; u_r_ is the associated vector of non-genetic random effects; g is the vector of additive genetic random effects with W the incidence matrix; and ε is the vector of residuals. It is assumed that g ~ *MVN*(0, Mσ_g_ ^2^), where σ_g_ ^2^ is the additive genetic variance and M is the genomic relationship matrix (GRM). For each trait in each age class, we ran this model twice: first with M being a GRM calculated from the 50K genotype data, and second with M being a GRM calculated from the imputed SNP genotypes. The GRMs were computed using DISSECT (Canela-Xandri et al. 2015) using VanRaden’s Method 2 GRM calculation, for which the genetic relationship between individuals i and j is computed as:

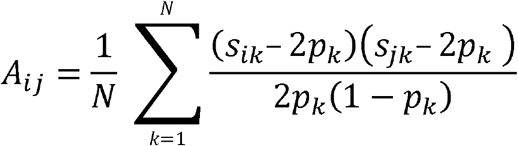

where s_ik_ is the number of copies of the reference allele for SNP k of the individual i, p_k_ is the frequency of the reference allele for the SNP k, and N is the number of SNPs (VanRaden 2008).

The narrow sense heritability was estimated by dividing the additive genetic variance (the variance explained by the GRM) by the total estimated phenotypic variance (the sum of the variance explained by the GRM and other fitted random effects after fitting fixed effects).

In adults, there are multiple records for August weight, foreleg length and hindleg length for the same individual due to individuals being caught across multiple years. For these traits we used a repeatability model in order that uncertainty was correctly propagated through all estimations (Mrode 2014). To implement a repeatability model in DISSECT, we edited the input files so that each measurement had its own row in the genotype and covariate files. Individual ID was replaced with a unique capture reference number, and individual permanent environment was fitted as a random effect (see Supplementary Methods for a more detailed explanation).

Sample sizes and total number of phenotypic measurements for all traits are shown in Table 1, with effects fitted in all models.

### Genome wide association analysis

Principal component analysis (PCA) using the GRM was performed prior to the genome-wide association analyses (GWAS) using DISSECT (Canela-Xandri et al. 2015) in order to examine the underlying population structure.

GWAS was also conducted using DISSECT (Canela-Xandri et al. 2015) using the following model:

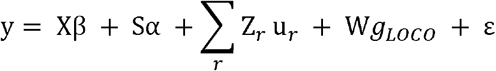

where y, X, β, Z_r_, u_r_, W and ∊ are the same as in the model for heritability estimation, s is the effect of the focal SNP, S is the design matrix linking individual records to the number of effect alleles for α, the estimated SNP effect (coded as 0, 1 or 2), and gLOCO is the vector of additive genetic random effects excluding the chromosome on which the focal SNP resides.

We fitted the same fixed and random effects for each trait and age class as for the heritability estimation (Table 1). To account for population structure, when testing SNPs on a given chromosome for association with the phenotype, a GRM calculated from the remaining autosomes (referred to as Leave One Chromosome Out GRM (gLOCO) (Yang et al. 2014)) was fitted. Input files for repeated-measure traits were reformatted as above. Our significance threshold was corrected for multiple testing using the SimpleM method (Gao et al. 2008), which accounts for linkage disequilibrium between markers in order to calculate the effective number of independent tests.

We estimated the variance explained by SNPs that passed the significance threshold using the equation

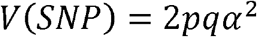

where p and q are the major and minor allele frequencies of the SNP, and α is the estimated SNP effect. We then calculated the proportion of additive genetic variance explained by each SNP by dividing by the total additive genetic variance estimated for that trait.

For any trait for which several SNPs in the same region were associated with variation in the trait and thus had strong support for at least one QTL in the region, we carried out conditional analysis to understand if the region could potentially harbour several independent QTL, or if further QTL could be uncovered elsewhere in the genome. To that aim, the genotypes of the SNP with the smallest association p value from each associated region (hereafter called the “top SNP”) were added to the GWAS model as a fixed covariate and removed from the GRMs and genotype data. The GWAS analysis was re-run accounting for those associations to try and reveal novel peaks either in the same regions or elsewhere in the genome.

### Genes in QTL regions

For each trait x SNP association, we investigated the genes within a 0.5Mb window either side of the top SNP to identify any genes which could be contributing to trait variation. We extracted a list of genes for each trait using the biomaRt package in R (Durinck et al. 2005; Durinck et al. 2009) from the OAR_v3.1 genome assembly and reviewed each gene against the NCBI Gene (Bethesda (MD): National Library of Medicine (US) 2004 – 2022) (including information from the Sheep Tissue Atlas (Jiang et al. 2014)), Animal QTLdb (Hu et al. 2022), and Ensembl (Howe et al. 2020) databases to examine function and expression annotations. When possible, we also compared with human and mouse orthologues due to the high level of annotation data available for these two species.

### Two-step GWAS analysis

GWAS has previously been performed on the adult traits in a smaller sample of Soay sheep using the 50K SNP data (Bérénos et al. 2015). The authors performed GWAS by first running a mixed model analysis, fitting fixed and random effects including whole-genome relatedness in the form of a GRM and, for repeated-measure traits, permanent environment. The residuals were then extracted and used as the phenotypic values for GWAS. For repeated measure traits, the mean residual value was used for each individual.

To investigate whether any novel SNP associations identified in this study were due to the increased sample size or due to the change in methodology, we also performed a two-step GWAS, focusing on adults only and using the 50K SNP data. Like Bérénos et al. (2015), we performed mixed model analyses using ASReml-R (Butler et al. 2017) and performed GWAS with the residuals as the trait phenotypes using DISSECT (Canela-Xandri et al. 2015). We used the Bonferroni correction calculated in Bérénos et al. (2015) to determine the significance threshold of 1.35e-6 for our two-step GWAS in order to compare with the previously published analysis.

## RESULTS

### Heritability estimation

#### Neonates

In neonates, the heritability of birth weight was 0.051 (S.E. 0.020) both when using the 50K SNPs to calculate relatedness, and when using the imputed SNPs (Figure 1, Supplementary Table 1). Given that both estimates are identical to 3 decimal places, there is no difference between the estimates.

**Figure 1.**
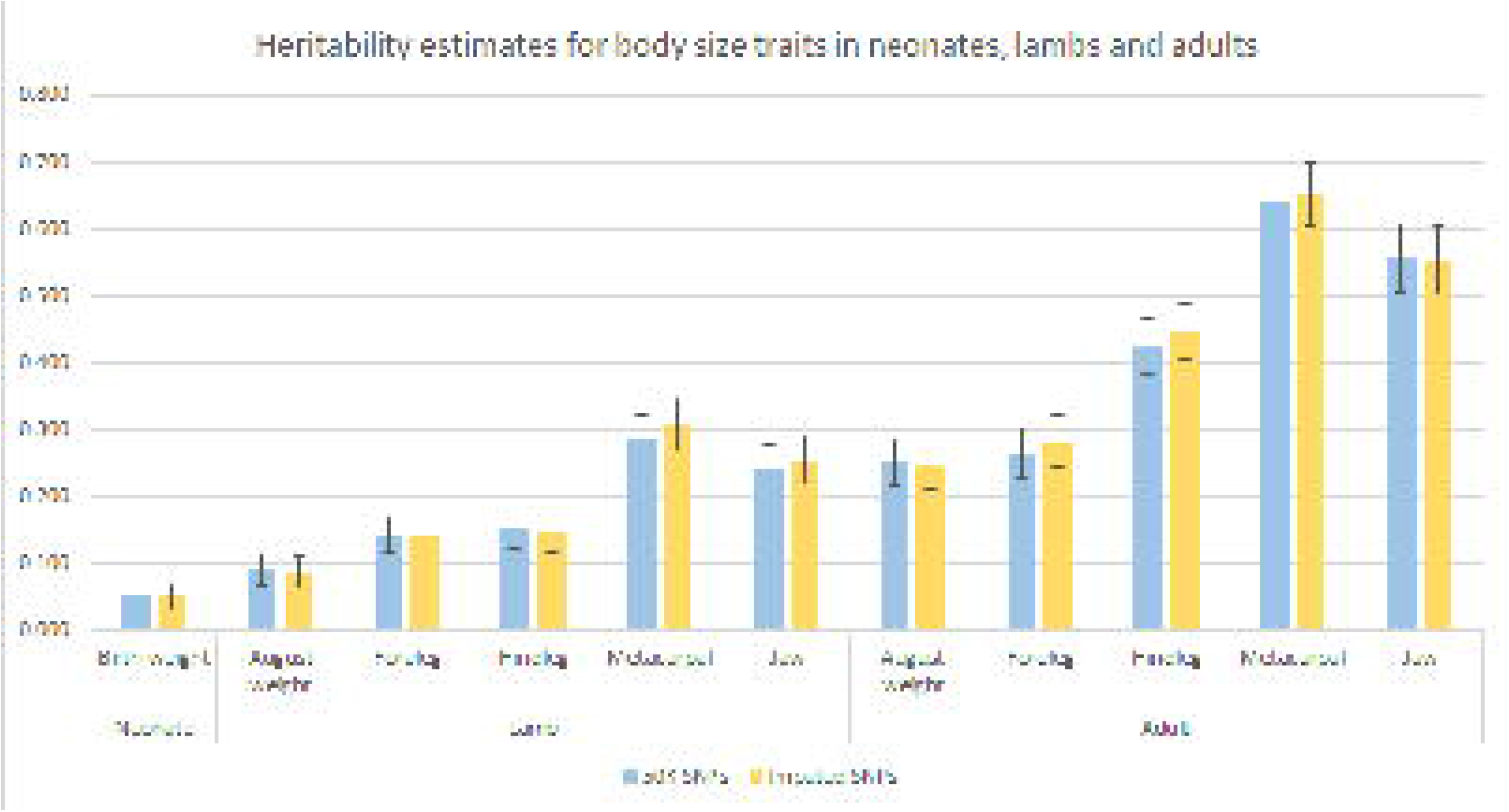
Estimates of heritability for body size traits in neonates, lambs, and adult Soay sheep when using a GRM calculated from the 50K SNP data (blue) compared with using a GRM calculated from the imputed SNP data (yellow). Error bars represent standard error estimates.

### Lambs

In lambs, the heritability estimates for the live August measures were lower than those for the post mortem measures (Figure 1, Supplementary Table 1). Across all the traits, heritability estimates were similar when using the 50K SNP data and the imputed SNP data, with the biggest difference being 0.024 for metacarpal length. For all traits, estimates were within one standard error of each other, indicating that the small differences in heritability estimates between the two SNP densities were not significant.

### Adults

As observed in lambs, heritability estimates for live measures in adults were lower than those of the post mortem measures. Across all traits, heritability estimates were higher in adults than in lambs. Estimates obtained using the 50K SNPs and using the imputed SNPs were similar and were within one standard error of each other (Figure 1, Supplementary Table 1), meaning that the imputed SNPs provided no additional information to partition the variation into genetic and environmental variance.

Estimates for all variance components are listed in Supplementary Table 1.

### GWAS

#### PCA

The top 20 principal components together explained 10.68% of the variance in the genetic data, with the top two principal components explaining 0.98% and 0.85% of the variance. We concluded that any population structure that was likely to affect the GWAS results would be corrected for by fitting a LOCO GRM in our GWAS model, as this will account for any structure caused by relationships.

### 50K SNP data

To correct for multiple testing, we calculated the effective number of tests to be 20082 using the SimpleM method (Gao et al. 2008), giving a genome-wide significance threshold of 2.49e^−06^ for the 50K SNP data.

For weight in neonates (birth weight), and lambs (August weight), no SNPs were found to have an association p value smaller than this threshold, suggesting that any variants that influence weight variation are either of small effect or were not tagged by SNPs in the 50K SNP data (Figure 2A, Supplementary Figure 1B and 1G). For adult August weight, three SNPs had a p value lower than the genome-wide significance threshold; one SNP on chromosome 6 and two SNPs on chromosome 9.

**Figure 2.**
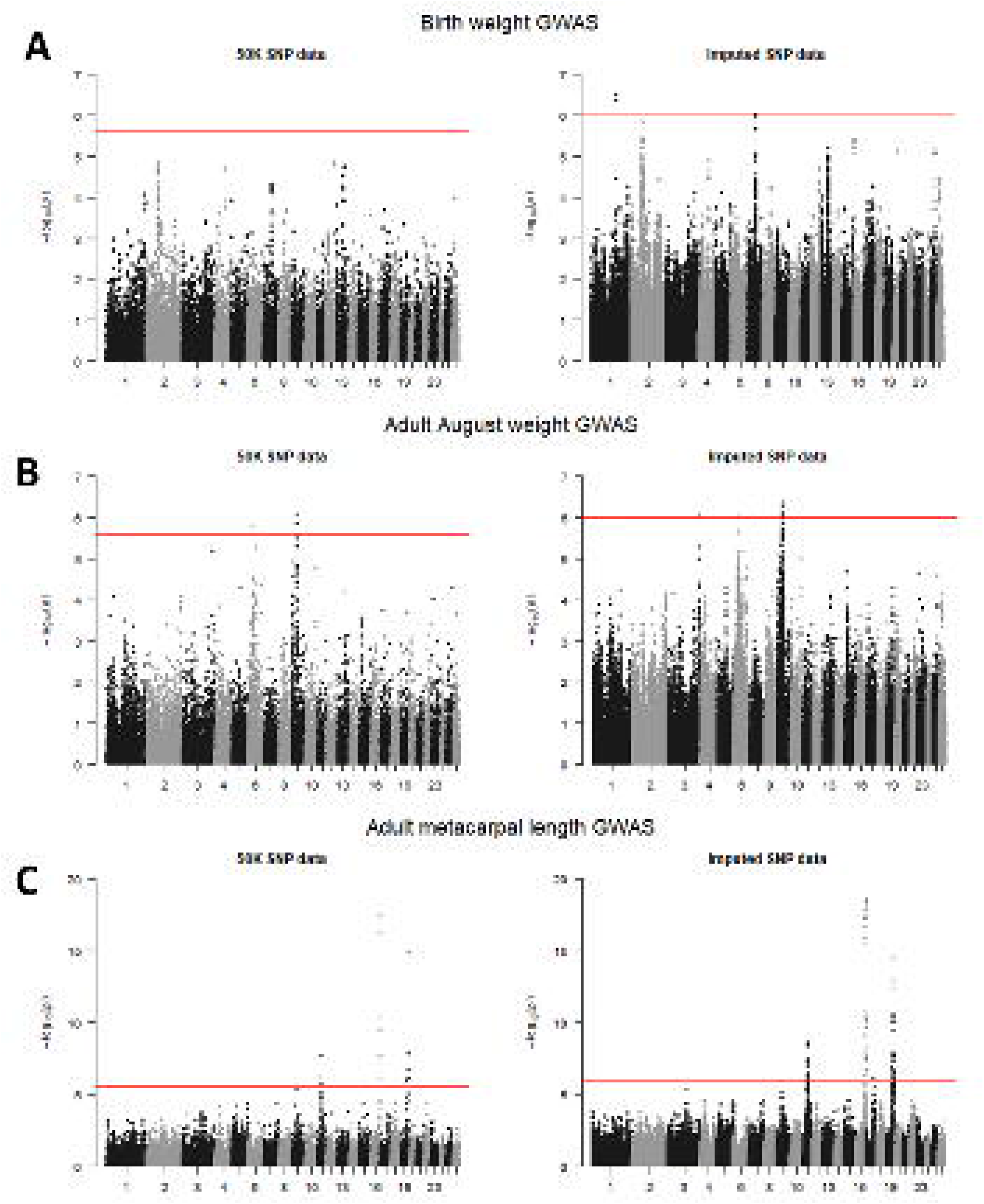
Manhattan plots for A) birth weight GWAS using 50K SNP data (left) and imputed SNP data (right); B) adult August weight GWAS using 50K SNP data (left) and imputed SNP data (right); and C) adult metacarpal length GWAS using 50K SNP data (left) and imputed SNP data (right). The red line represents the significance threshold (2.49e^−06^ for the 50K SNP data and 1.03e^−06^ for the imputed SNP data) – any SNPs above this threshold are considered to be significantly associated with variation in their respective traits.

For all three leg length measures in lambs, we found associations with the same region on chromosome 16. SNP s23172.1 was the SNP with the lowest p value for lamb foreleg and hindleg, explaining 0.52% and 0.69% of the genetic variance for each trait respectively (Supplementary Table 2, Supplementary Figure 1C and 1D). For lamb metacarpal, SNP 22142.1 in the same chromosome 16 region had the lowest p value and explained 0.97% of the genetic variance. There was also a single SNP on chromosome 3 (OAR3_100483326.1) and a cluster of SNPs on chromosome 19 that had p values smaller than the genome-wide significance threshold and were associated with variation in lamb metacarpal length, with the SNP with the lowest p value from each region explaining 2.08% and 2.40% of the genetic variance respectively (Supplementary Table 2, Supplementary Figure 1E).

**Table 2.** Potential candidate genes for future analyses. From left to right: gene name, Ensembl gene ID, chromosome, associated trait, and evidence for association in other species.

| Gene Name | Ensembl Gene ID | Chr | Associated trait | Effects in other species |
| --- | --- | --- | --- | --- |
| Cytochrome P450 26B1 | ENSOARG00000011582 | 3 | Lamb metacarpal length | Associated with skeletal abnormalities in humans and zebrafish (Laue et al. 2011), knockouts produce reduced limbs in mice (Yashiro et al. 2004). |
| EXOC6B | ENSOARG00000011607 | 3 | Lamb metacarpal length | Associated with spondyloepimetaphyseal dysplasia (resulting in short stature) in humans (Campos-Xavier et al. 2018). |
| FAM13A | ENSOARG00000018727 | 6 | Adult August weight | Modulates body fat distribution and adipocyte function in humans and mice (Fathzadeh et al. 2020) as well as adipose insulin signalling in mice (Wardhana et al. 2018), also linked with obesity in mice (Tang et al. 2019) |
| ONECUT1 | ENSOARG00000020928 | 7 | Birth weight | Associated with birth weight in humans (Warrington et al. 2019). |
| IFT43 | ENSOARG00000002065 | 7 | Adult foreleg length | Associated with Sensenbrenner syndrome (resulting in growth retardation and dwarfism due to femoral and humeral limb shortening) in humans (Arts et al. 2011). |
| PENK | ENSOARG00000020184 | 9 | Adult August weight | PENK knock-out mice found to have diminished food motivation, lower baseline body weight and attenuated weight gain (Mendez et al. 2015) |
| PTH1R | ENSOARG00000006638 | 19 | Lamb metacarpal length, adult foreleg length, adult hindleg length, adult metacarpal length | Involved in osteoblast development in mice (Qiu et al. 2015), associated with skeletal disorders such as EKNS (Duchatelet et al. 2005), JMC and BLC (Schipani and Provot 2003) in humans. |
| LTF | ENSOARG00000008620 | 19 | Lamb metacarpal length, adult metacarpal length | Human LTF associated with increased bone growth when injected into piglets (Li et al. 2018), found to stimulate osteoblast proliferation (Cornish and Naot 2010). High expression levels in human bone marrow (Fagerberg et al. 2014). |
| BMP6 | ENSOARG00000017264 | 20 | Adult jaw length | Involved in bone development and expressed in the jaw bone in mice (Oralová et al. 2014). |

The two regions on chromosomes 16 and 19 that were associated with lamb metacarpal length variation were also significantly associated with all three leg length measures in adults, with SNP s22142.1 on chromosome 16 and SNP s74894.1 on chromosome 19 respectively explaining 0.80% and 2.04% of the genetic variation in adult foreleg, 0.88% and 1.32% of the genetic variation in adult hindleg, and 0.55% and 2.02% of the genetic variation in adult metacarpal length. There were other regions of the genome also associated with variation in the adult leg length traits; a region on chromosome 11 was significant across all three adult leg length traits, with the most significant SNP explaining 2.35%, 2.25% and 1.13% of the genetic variance in adult foreleg, hindleg and metacarpal respectively (Figure 2B, Supplementary Table 2, Supplementary Figure 1H and 1J). For adult foreleg, a SNP on chromosome 7 and two on chromosome 9 were also associated, with the most significant SNPs in each region explaining 1.31% and 2.99% of the genetic variance respectively for this trait (Supplementary Table 2, Supplementary Figure 1H).

In lambs, there were no associations with jaw length found (Supplementary Figure 1F). In adults, a SNP on chromosome 20 was associated with jaw length variation, explaining 2.05% of the genetic variance for this trait (Supplementary Table 2, Supplementary Figure 1K).

In total, we identified 85 SNP-trait associations with 39 unique SNPs.

### Imputed data

Using the SimpleM method (Gao et al. 2008), we calculated the number of effective tests to be 48635, giving a genome-wide significance threshold of 1.03e^−06^.

When performing GWAS using the imputed SNP data, we were able to recover significant SNPs in the same locations for all traits as those we found using the 50K SNP data. Of the 85 SNP-trait associations that we identified with the 50K SNP data, 81 were significant using the imputed SNP data – the remaining four SNPs were no longer significant due to the increased multiple testing burden (which leads to a more stringent significance threshold) between the 50K SNP data and the imputed SNP data (2.49e^−06^ and 1.03e^−06^ respectively).

We also identified 795 new SNP-trait associations using the imputed SNP data with 425 unique SNPs (Supplementary Table 2). The majority of new associations were in the same regions as the SNPs identified using the 50K SNP data, but we also found new associations: four SNPs on chromosome 1 and three SNP on chromosome 7 was associated with birth weight (Figure 2A, Supplementary Table 2), one SNP on chromosome 3 was associated with adult August weight (Figure 2B, Supplementary Table 2), and one SNP on chromosome 17 was associated with adult metacarpal length (Figure 2C, Supplementary Table 2).

Manhattan and QQ plots for all traits can be found in Supplementary Figure 1.

### Conditional analysis

We performed conditional analysis on all three leg length traits in both lambs and adults, as well as on birth weight, adult August weight adult jaw length (See Supplementary Table 2 for all SNPs that were fitted for each trait). For all of these traits we performed the conditional analysis using both the 50K SNP data and the imputed SNP data, with the exception of birth weight, which did not have any significant SNP associations using the 50K data.

Six of the nine traits we performed conditional analysis on had significant SNPs after fitting the SNPs with the lowest p value, however for four of these traits (lamb metacarpal length, adult August weight, foreleg length and hindleg length), these were SNPs that were also significant in our original GWAS analysis but were not fitted in the conditional analysis due to being the only SNP that was significantly associated with the trait in that region (Supplementary Table 3). The remaining two traits (birth weight and adult jaw length) both had a new association, both of which were on chromosome 2. For birth weight, nine SNPs had p values lower than the genome-wide significance threshold, all around ~81Mb (Figure 3A, Supplementary Table 3). For adult jaw length, only one SNP had a lower p value than the genome-wide significance threshold, at position 137,162,126 (Figure 3B, Supplementary Table 3).

**Figure 3.**
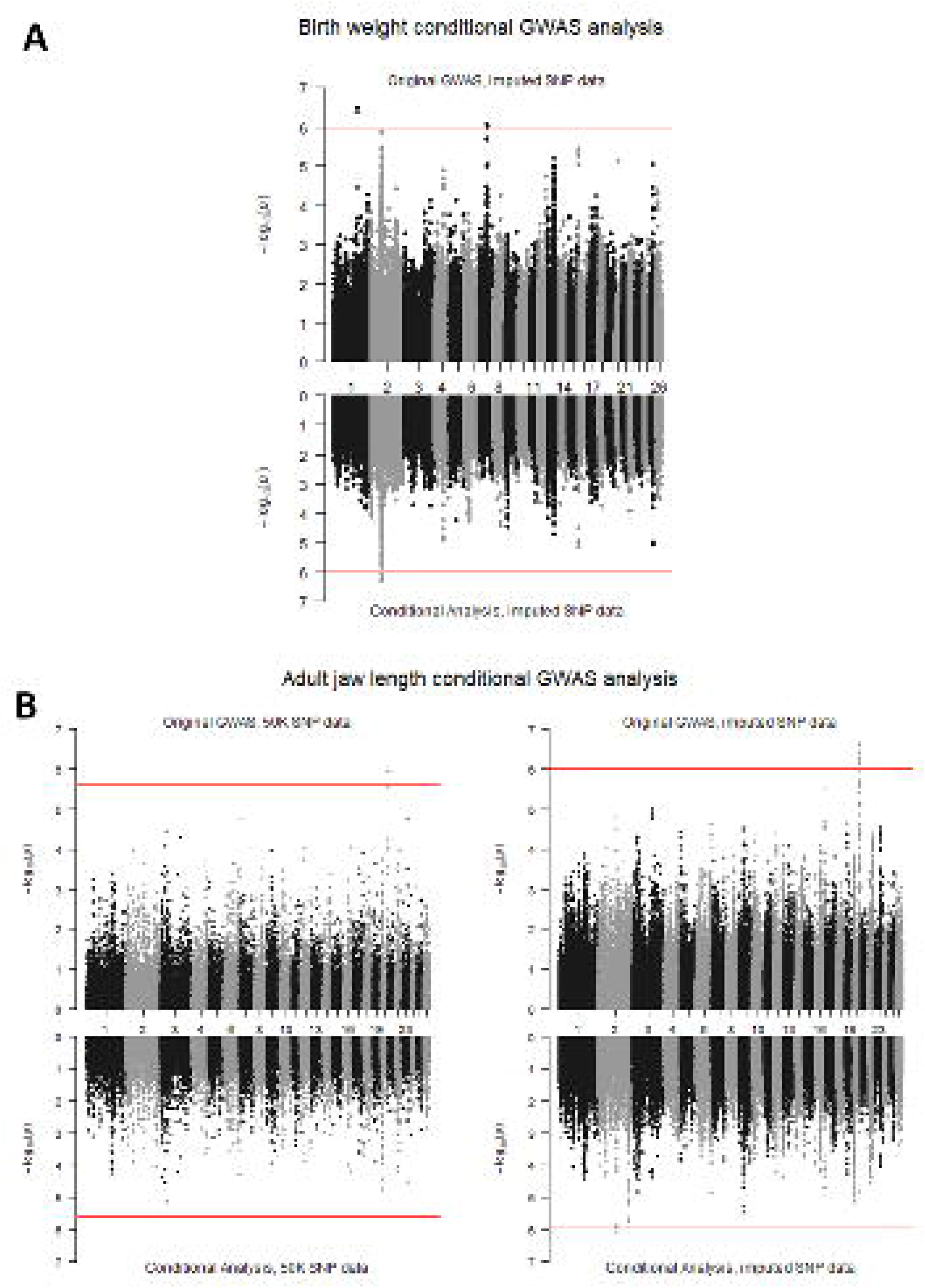
Miami plots for A) birth weight using imputed SNP data (top) and birth weight conditional analysis using imputed SNP data (bottom); and B) adult jaw length GWAS using 50K SNP data (top left) and imputed SNP data (top right), adult jaw length conditional analysis using 50K SNP data (bottom left) and imputed SNP data (bottom right). The red line represents the significance threshold (2.49e^−06^ for the 50K SNP data and 1.03e^−06^ for the imputed SNP data) – any SNPs above this threshold are considered to be significantly associated with variation in their respective traits.

### Genes in QTL regions

Given that all of the region-trait associations that were found to be significant with the 50K SNP data were also significant with the imputed SNP data, we chose to focus on top SNPs in the imputed dataset, including the two regions that were only significantly associated when performing the conditional analysis (See Supplementary Table 2 and 3 for the list of SNPs, and Supplementary Table 4 for the list of genes).

We found a total of 179 genes in the regions around the SNPs associated with our traits. 56 of these genes were unannotated in the current sheep genome build, and of those that were annotated, three did not have a listed mouse homologue and a further six had neither a mouse nor a human homologue. Of the genes that did have annotation and homologue data, we found nine that are associated with similar traits to our focal traits in humans and mice, suggesting that they may be contributing to the genetic variation of our traits (Table 2). However, without intimate knowledge of the genes surrounding the focal SNPs, it is likely that there are other genes that are also contributing. It is also worth noting that variation in coding regions of genes in proximity to SNPs we identified as being associated with our traits may not be responsible for variation in the phenotypes, but instead the causal variants may lie in regulatory sequences that modulate expression of either these or other genes.

We also compared our GWAS results with QTL from Animal QTLdb (Hu et al. 2022). We found that the region on chromosome 6 that we found to be associated with adult August weight overlaps with a region previously found to be associated with carcass weight and final body weight in an (Awassi x Merino) x Merino backcross population (Cavanagh et al. 2010) and is ~0.5Mb upstream of a 2.5Mb region that has also previously been associated with body weight in a population of Australian Merino sheep (Al-Mamun et al. 2015). In addition, the region on chromosome 9 we found to be associated with adult August weight is 1Mb upstream of a region previously found to be associated with yearling live weight in a population of Chinese Merino sheep.

Chromosome 6 has previously been associated with adult body weight using linkage analysis in a smaller sample of Soays (Beraldi et al. 2007), however the microsatellite markers flanking the associated region are not located close to the region we identified.

### Two-step GWAS

Across all 5 traits when using the two-step approach, we recovered the SNP-trait associations identified by Bérénos et al. (2015). However, we were unable to recover any of the novel SNP-trait associations we found when performing our single-step GWAS on the 50K SNP data, with the exception of the association between chromosome 16 and adult foreleg (though Bérénos et al. noted that SNPs in this region approached significance in their analysis). Despite the genome-wide significance threshold used by Bérénos et al. (2015) being more stringent than the significance threshold we calculated using the SimpleM method for the 50K SNP data, no additional associations are recovered when using our less stringent threshold

Our QQ plots using the two-step method also matched the QQ plots of Bérénos et al. (2015). In both, the observed p values were higher than the expected p values, causing the majority of points in the plots to fall below the x=y line (Supplementary Figure 3).

## DISCUSSION

### Heritability

Our results corroborate findings from previous studies in the Soay sheep. For instance, we showed that all five body size traits we studied in Soay sheep are influenced by genetic variation in the population (Bérénos et al. 2014), and that post mortem measures have higher heritability estimates than live measures. We also found that leg measures have higher heritability than weight (Wilson et al. 2006; Beraldi et al. 2007; Bérénos et al. 2014), and that heritability estimates increase with age (Wilson et al. 2006; Bérénos et al. 2014).

The heritability estimates for the 50K data were very similar to those estimated using a GRM based on the 50K data in a smaller sample of the same population of sheep by Bérénos et al. (2014), with estimates for the same trait falling within one standard error of each other. The biggest difference was in adult metacarpal length with a heritability difference of 0.05 (estimates were 0.644 (0.047) and 0.594 (0.047) for our and Bérénos et al.’s results respectively). Given that we used the same models as Bérénos et al., it is likely that the small differences between heritability estimates for each trait is due to our increased sample sizes.

Comparing the heritabilities estimated using the imputed SNP data against the estimates using the 50K SNP data, we found little difference between the two SNP densities in any traits in any age class. The additional genotypes at the imputed SNPs do not give any additional information on additive genetic variation for these traits. This result is not surprising given the previous rarefaction analysis showing that the heritability of these body size traits in adults asymptoted when about half the 50K SNP data was used (Bérénos et al. 2014). There is high LD between nearby SNPs in the Soay sheep genome, which suggests that most, if not all, of the causal variants tagged by the imputed SNP data may have already been tagged by the 50K SNPs. The high LD was reflected when calculating GWAS significance thresholds – whilst the number of SNPs between the 50K SNP data and the imputed SNP data increased by a factor of ten, the number of effective tests only doubled (39K SNPs, 20082 effective tests and 401K SNPs and 48635 effective tests respectively).

For some of the traits we have analysed there is still a difference in heritability estimated using SNP data versus heritability estimated using pedigree – for example, the highest SNP-based heritability estimate for lamb metacarpal length (the estimate using the imputed SNP data) gave an estimate 59% of Bérénos et al.’s pedigree-based estimate (Bérénos et al. 2014). Given that our SNP-based heritability estimates were similar when using the 50K SNP data as when using the imputed SNP data, and the results of Bérénos et al.’s rarefaction analysis (Bérénos et al. 2014), we believe it is unlikely that increasing the density of genotyped SNPs that are common in the population will increase heritability estimates of these traits. It is possible instead that the difference in heritability estimates obtained from pedigree and genomic data is due to rare familial variants that do not segregate widely in the population, as well as due to dominance and epistasis.

## GWAS

Body size traits have been the focus of many kinds of analyses in Soay sheep (Beraldi et al. 2007; Ozgul et al. 2009; Bérénos et al. 2014; Bérénos et al. 2015; Pemberton et al. 2017; Regan et al. 2017; Ashraf et al. 2021), and several SNP associations have already been identified for these traits. A 2015 study aiming to find SNP-trait associations identified QTL for adult leg length measures on chromosomes 16 and 19 (s23172.1 and s74894.1 respectively) (Bérénos et al. 2015). A more recent study comparing genomic prediction methods in Soays using the 50K SNP data identified s48811.1 on chromosome 7 and s50107.1 on chromosome 9 as having a probability higher than 0.9 of having a non-zero effect on adult foreleg length in addition to the previously discovered regions on chromosomes 16 and 19 (Ashraf et al. 2021). We were able to identify all four of these associations in our GWAS, alongside associations that have not previously been identified in this population. Use of the imputed SNP data allowed us to discover four more associations with loci that were not genotyped in the 50K SNP data, indicating that future identification of polymorphisms influencing trait variation in the Soay sheep will benefit from using the imputed data.

Performing a two-step analysis confirmed that the novel SNP-trait associations we were able to identify using the 50K SNP data were due to fitting the fixed and random effects for each trait simultaneously, performing GWAS in a single step, rather than due to having a larger sample size. Given the increase in SNP-trait associations when using the single-step methodology, and with the availability of software like DISSECT which is able to fit fixed and random effects whilst performing GWAS, two-step GWAS should be avoided. As we have shown, although DISSECT does not currently have the option to automatically run a repeated measures GWAS, it is possible to modify input files to allow for repeated measures to be appropriately modelled.

The imputed SNP data revealed SNP-trait associations in four regions of the genome that were not discovered using the 50K SNP data; a region on chromosome 1 and a region on chromosome 7 and birth weight, a region on chromosome 3 and adult August weight, and a region on chromosome 17 and adult metacarpal length. (Supplementary Table 2). When examining the Manhattan plot for the 50K data for each trait (Figure 2A, 2B and 2C, Supplementary Table 2) it is clear that, with the exception of the region on chromosome 1 associated with birth weight, there was a small cluster of SNPs just under the significance threshold in the 50K analyses. The additional (imputed) SNPs may have matched the allele frequency of the underlying causal variants more accurately, resulting in a smaller association p value.

We performed conditional analysis on all three leg length traits in both lambs and adults, as well as on birth weight (only using the imputed SNP data), adult August weight and adult jaw length. For each trait, we simultaneously fitted the genotype for the SNP with the lowest p value for any chromosome that had at least two SNPs found to be associated with the trait (see Supplementary Table 2 for a list of SNPs fitted for each trait). We found that all of the SNPs that were significant in the GWAS analysis were no longer significant in the conditional analysis when a significant SNP on the same chromosome was fitted (Figure 3, Supplementary Table 3). We suggest that any future work looking to pinpoint the exact location of the genetic variants affecting body size traits in Soay sheep primarily focus on the regions around the SNPs listed in Supplementary Table 2.

We identified 179 genes within 0.5Mb of the top SNPs for each trait (Supplementary Table 4), and of these genes, we found nine that are potential candidate genes for further analyses due to their association with similar traits in other species. Two of these genes (*CYP26B1* and *EXOC6B*) are associated with the same trait (lamb metacarpal length) and are located within 0.5Mb of the same top SNP on chromosome 3. *CYP26B1* is associated with skeletal abnormalities in humans and zebrafish (Laue et al. 2011) and *CYP26B1* knockouts produced reduced limbs in mice (Yashiro et al. 2004), whilst *EXOC6B* is associated with spondyloepimetaphyseal dysplasia in humans which symptoms include skeletal malformations affecting the long bones of the limbs and short stature (Campos-Xavier et al. 2018).

*PTH1R* is located in the region on chromosome 19 associated with lamb metacarpal length and all three adult leg traits, and has been found to be involved in osteoblast development in mice (Qiu et al. 2015), and is associated with skeletal disorders such as Eiken’s syndrome (Duchatelet et al. 2005), Jansen’s metaphyseal chondrodysplasia and Blomstrand’s lethal chondrodysplasia (Schipani and Provot 2003). As the top SNPs on chromosome 19 for both lamb and adult metacarpal are in a slightly different position to the top SNP for the live measures, *LTF* is in the region associated with both lamb and adult metacarpal length on chromosome 19, and has been found to increase bone growth when injected into piglets (Li et al. 2018), stimulate osteoblast proliferation in humans (Cornish and Naot 2010), and has high expression levels in human bone marrow (Fagerberg et al. 2014).

*BMP6*, located in the region on chromosome 20 associated with adult jaw length, has also been found to be involved in bone development and expression in mice jaw bone (Oralová et al. 2014).

We found three potential candidate genes associated with weight. *FAM13A* in the region on chromosome 6 associated with adult August weight has been found to modulate body fat distribution and adipocyte function in humans and mice (Fathzadeh et al. 2020), affects adipose insulin signalling in mice (Wardhana et al. 2018), and is associated with obesity also in mice (Tang et al. 2019). *PENK*, located in the region on chromosome 9 associated with adult August weight has been found to cause diminished food motivation, lower baseline body weight and attenuated weight gain when knocked out in mice (Mendez et al. 2015). *ONECUT1*, located in the region on chromosome 7 associated with birth weight has been found to be associated with birth weight in humans (Warrington et al. 2019). Unlike the other potential candidate gene-trait associations we found, this association was only found when performing GWAS with the imputed SNP data.

Although we have discovered new SNP-trait associations, it is likely that there are still causative variants that remain undetected. GWAS lacks power to detect rare causative variants and variants with very small effect sizes (Yang et al. 2010). Also, GWAS power drops when the same amount of phenotypic variation is a consequence of multiple variants in the same region as opposed to a single variant (Nagamine et al. 2012). Regional mapping methods have been developed that partition trait variance into regions by simultaneously fitting a whole genome and a regional GRM, with the regions either being defined as fixed SNP windows (Nagamine et al. 2012) or haplotype blocks (Shirali et al. 2018). Such methodologies have the potential to identify regions of the genome that contain variants associated with a trait that are unable to be identified by GWAS either due to being rare, or individually having small effects on trait variation. Genomic prediction, which simultaneously estimates all marker effects drawn from multiple distributions, can also be used to study the genetic architecture of traits by estimating the posterior inclusion probability of a SNP having a non-zero effect on a trait. Genomic prediction has already been used on adult body size traits in Soays, and has identified several of the SNPs we identified through our GWAS approach (Ashraf et al. 2021). Ultimately, we believe that it is important to use a variety of methodologies when studying the genetic architecture of complex traits, as different analyses have different strengths and may be able to identify different QTL.Across all traits for all age classes, the QQ plots showed deviation from the expected distribution of test statistics under the null hypothesis (x=y line) for a wide range of test statistics, including low values, which may be either due to underlying population structure not accounted for by the GRMs or due to trait architecture. The first 20 genomic principal components accounted for only 10.68% of the variance in the genetic data, and repeating the GWAS analysis fitting these first 20 genomic principal components in addition to the GRM did not change the p values of the SNPs nor the QQ plots (data not shown). This shows that the principal components in this case were not useful in adjusting for population structure in the presence of the GRM, and that population structure is not likely to cause any issues in our analyses, especially after fitting a LOCO GRM.

In order to have sufficient power to detect associations between markers and a trait of interest, GWAS primarily requires two factors: i) a very high density of genotyped SNPs, and ii) a large number of individuals that have been genotyped and phenotyped (Santure and Garant 2018). For intensively studied organisms, both are achievable; such populations tend to have more individuals accessible to collect data from, high density genotyping can be done at a lower cost due to higher demand, and, as in humans, data from different populations can be combined to create larger sample sizes. GWA studies of humans are the most obvious example of this; studies often have study populations made up of hundreds of thousands of individuals and human SNP chips commonly genotype hundreds of thousands of variants (for example, see Wood et al. 2014; Ishigaki et al. 2020; Wu et al. 2021). In comparison, wild study population samples are much smaller – often struggling to reach one thousand individuals – and the number of SNPs genotyped is much lower (for example, see Silva et al. 2017; Malenfant et al. 2018; Perrier et al. 2018). Analyses of wild populations therefore generally lack the power of more intensively studied study organisms. Here, we have increased power by increasing the number of genotyped markers via imputation. Despite high LD in the Soay sheep population, use of imputed data has allowed us to identify four new SNP-trait associations, including an association with birth weight, which had yet to be associated with any QTL in the Soay population. We have therefore shown that for a given sample size, more information can be obtained by increasing the density of markers for those individuals have been phenotyped. We suggest that, where possible, analyses of wild populations impute SNP data in order to increase power and obtain results that may otherwise remain undiscovered.

In our population, we were able to identify new SNP-trait associations when performing GWAS with the higher density imputed dataset that were not found with the lower density non-imputed data – one of which allowed for the identification of a potential candidate gene for that trait. We have shown that analyses performed with imputed data, such as heritability estimation and GWAS, can still pick up additive genetic variation that would have been identified with the non-imputed data. We suggest that, where possible, analyses of wild populations impute SNP data in order to increase power and obtain associations that may otherwise remain undiscovered. In addition, we demonstrated the benefits of fitting fixed and random effects during GWAS instead of pre-correcting and performing GWAS on the residuals. We therefore recommend that researchers use this “one-step” method when performing GWAS on wild populations.

## Data Availability Statement

All scripts and data can be found at https://github.com/CaelinnJames/Impact_of_SNPDensity_on_Soay_Sheep

## Acknowledgments

We thank the National Trust for Scotland and Scottish Natural Heritage for permission to work on St Kilda and QinetiQ and Eurest for logistics and other support on the island. We also thank all those who have been involved in the long-term project, including those who helped with field work on the island. We thank the Wellcome Trust Clinical Research Facility Genetics Core in Edinburgh for SNP genotyping.

