## Supplementary Figure 1 for "The impact of SNP density on quantitative genetic analyses of body size traits in a wild population of Soay sheep"

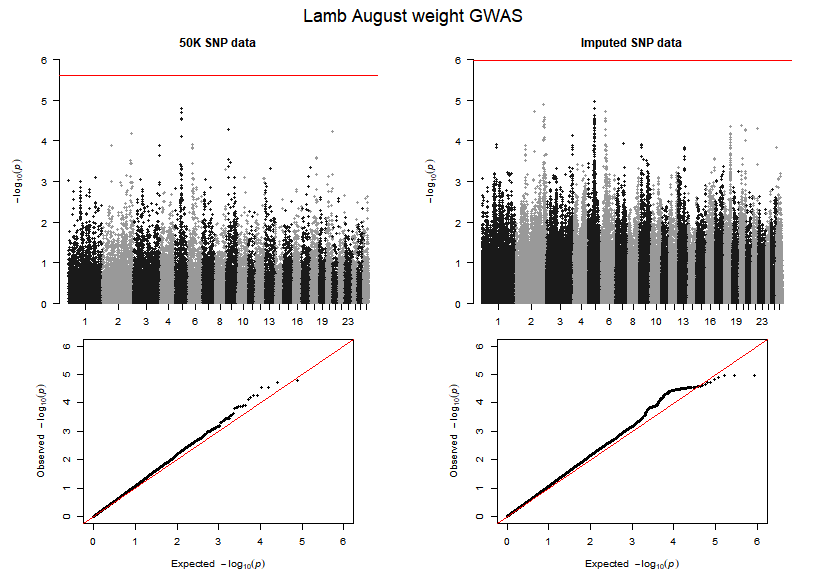

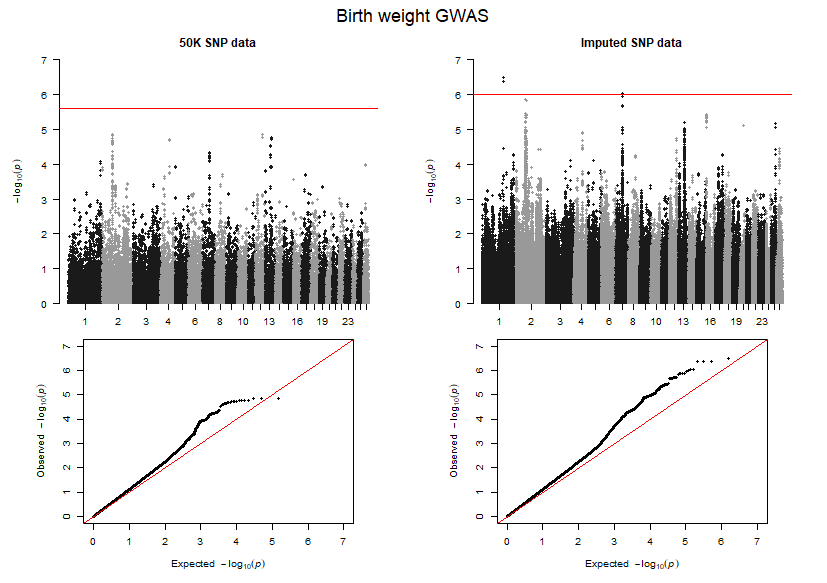

**A**

**B**

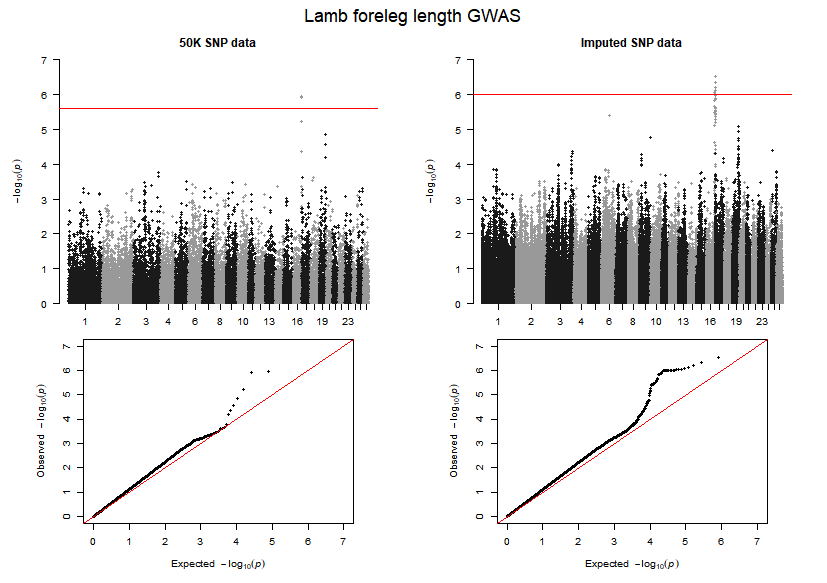

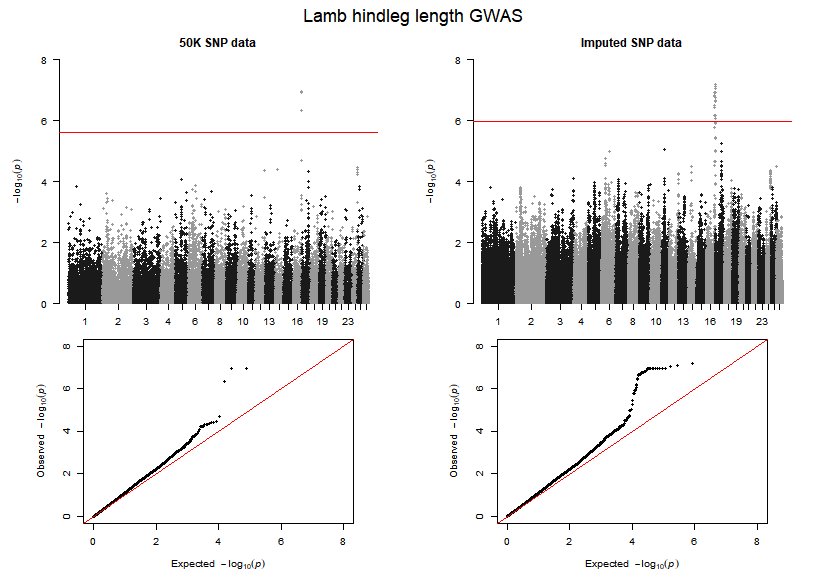

**C**

**D**

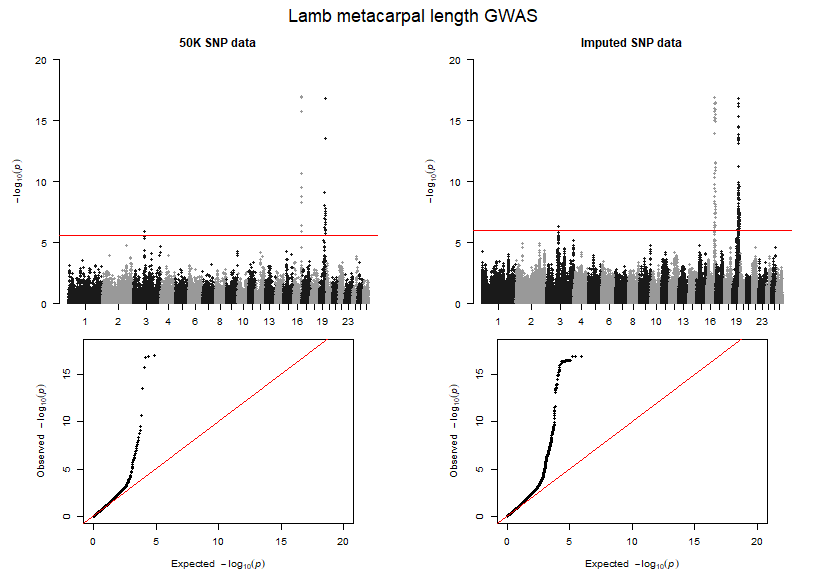

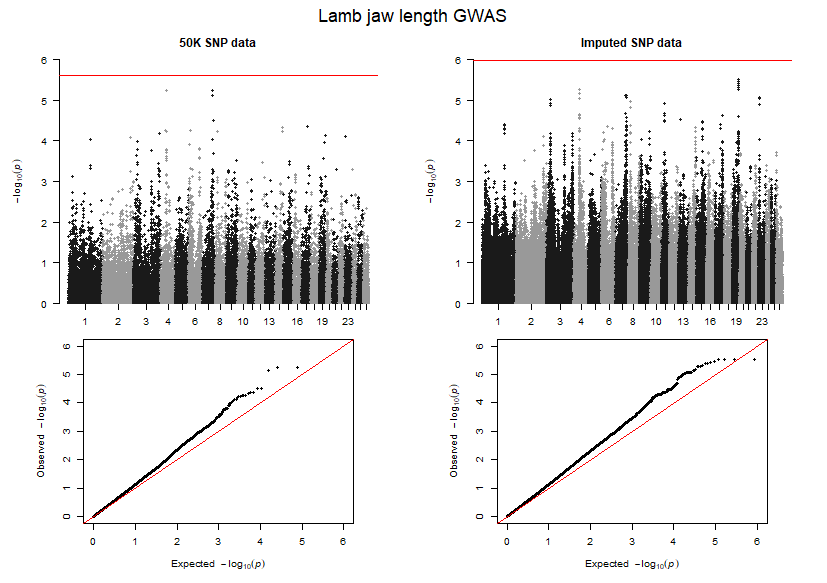

**E**

**F**

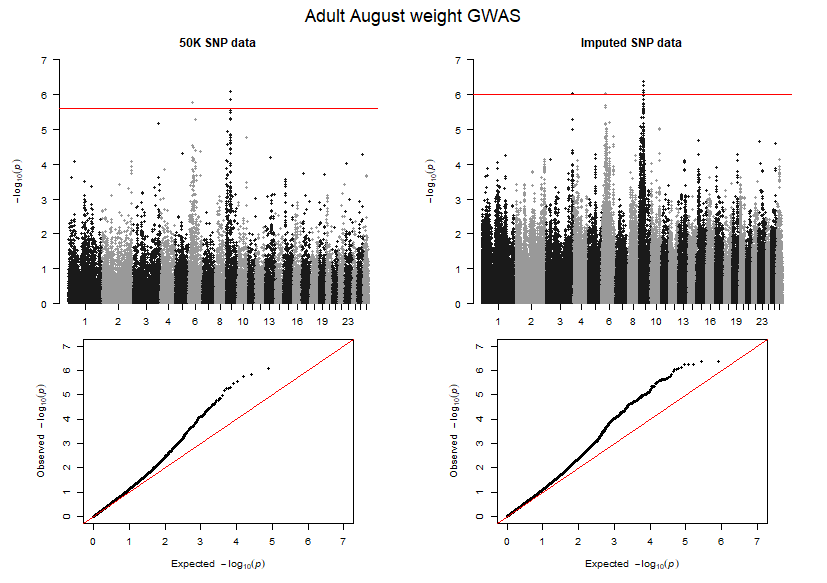

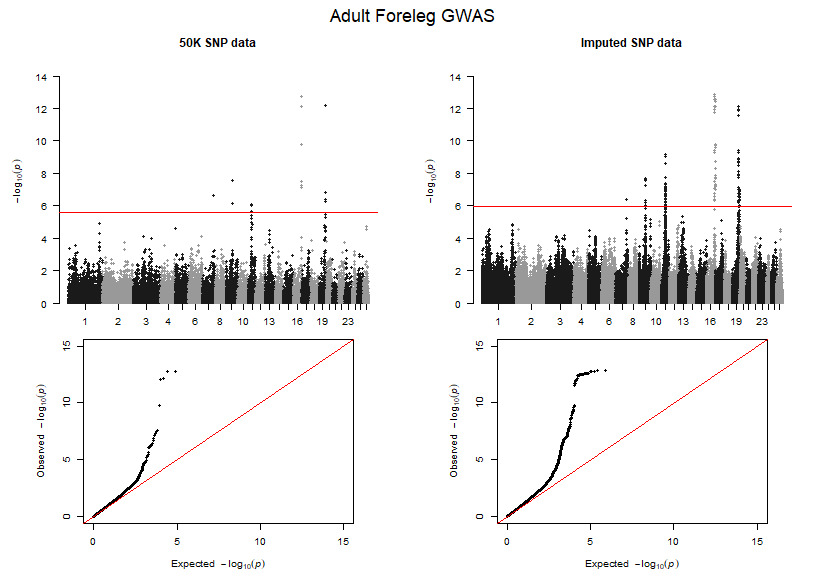

**G**

**H**

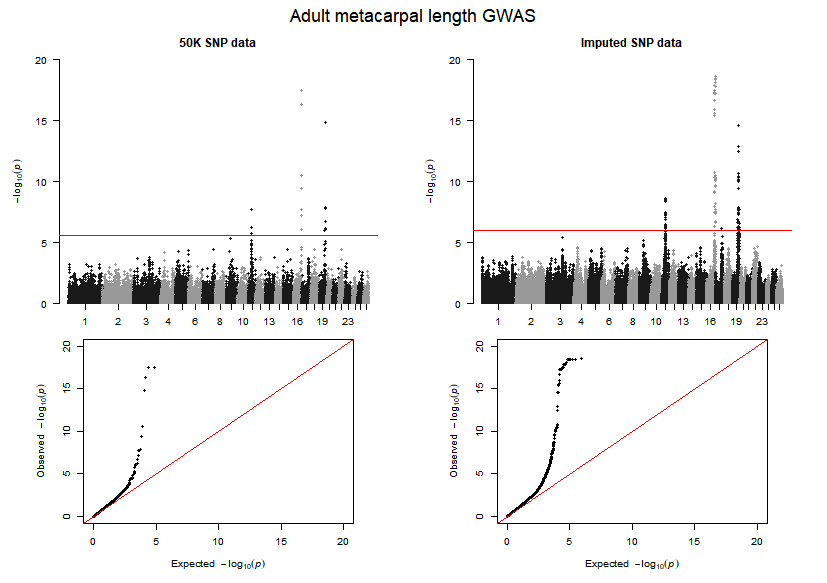

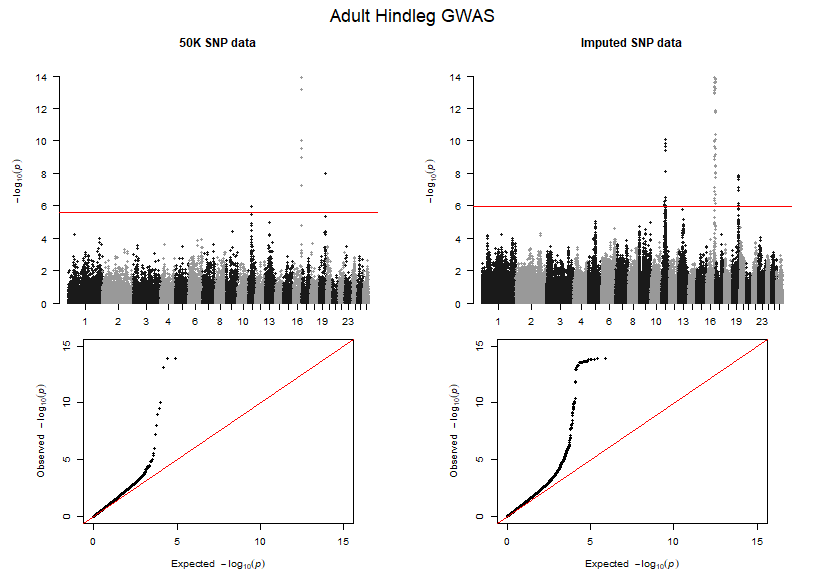

**I**

**J**

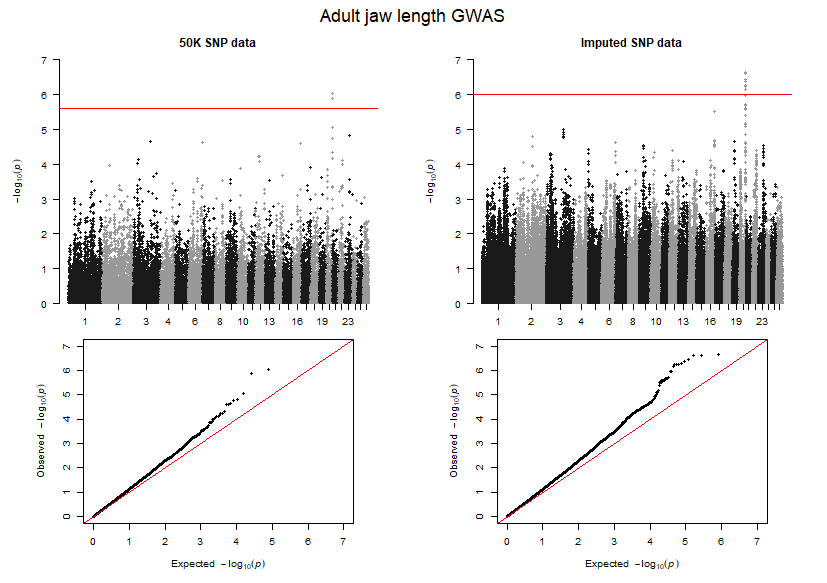

**K**

**Supplementary Figure 1** Manhattan and QQ plots using the 50K SNP data (left) and the imputed SNP data (right) for all traits and age classes. **A)** birth weight, **B)** lamb August weight, **C)** lamb foreleg length, **D)** lamb hindleg length, **E)** lamb metacarpal length, **F)** lamb jaw length, **G)** adult August weight, **H)** adult foreleg length, **I)** adult hindleg length, **J)** adult metacarpal length, and **K)** adult jaw length. In the Manhattan plots, the red line represents the significance threshold (2.49e^-06^ for the 50K SNP data and 1.03e^-06^ for the imputed SNP data) – any SNPs above this threshold are considered to be significantly associated with variation in their respective traits. In the QQ plots, the red line is the X=Y line.
