## Supplementary Figure 2 for "The impact of SNP density on quantitative genetic analyses of body size traits in a wild population of Soay sheep"

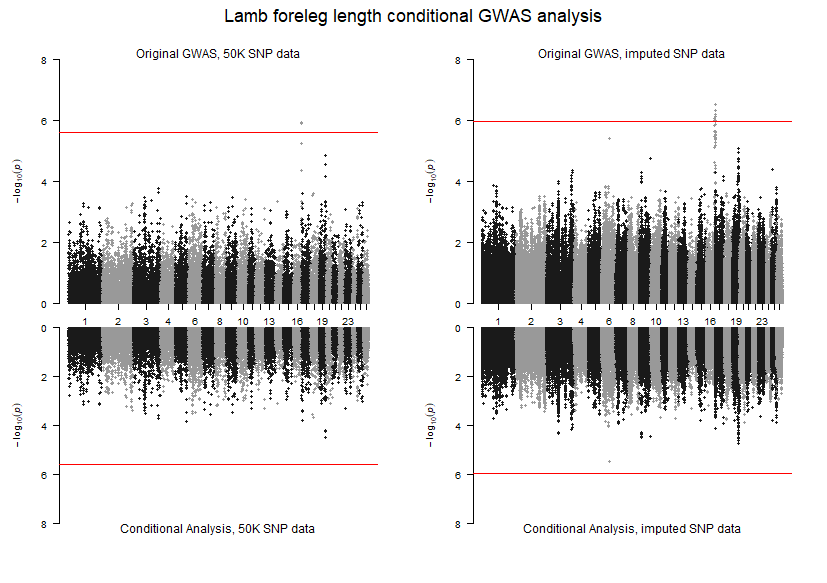

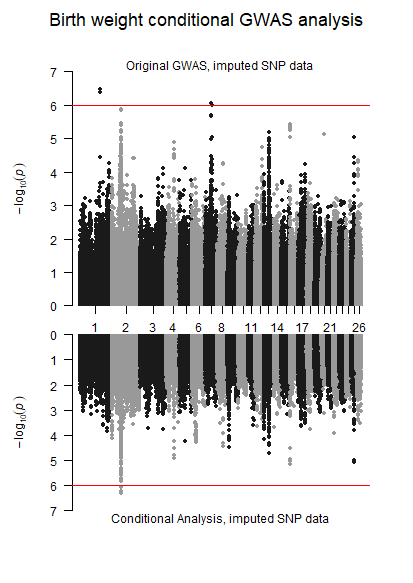

**A**

**B**

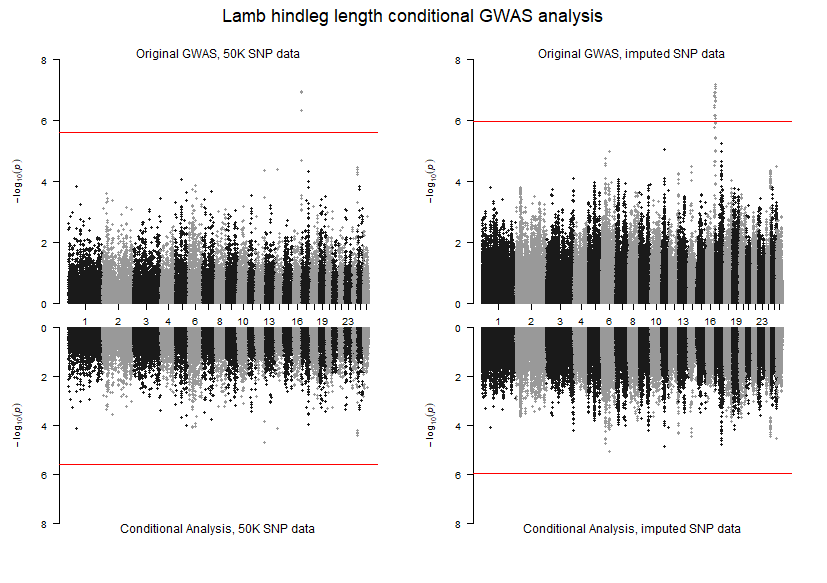

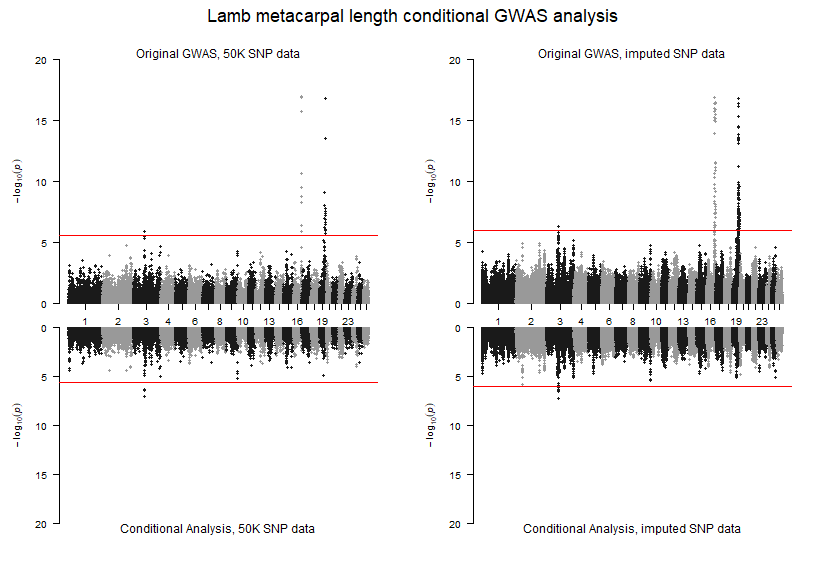

**C**

**D**

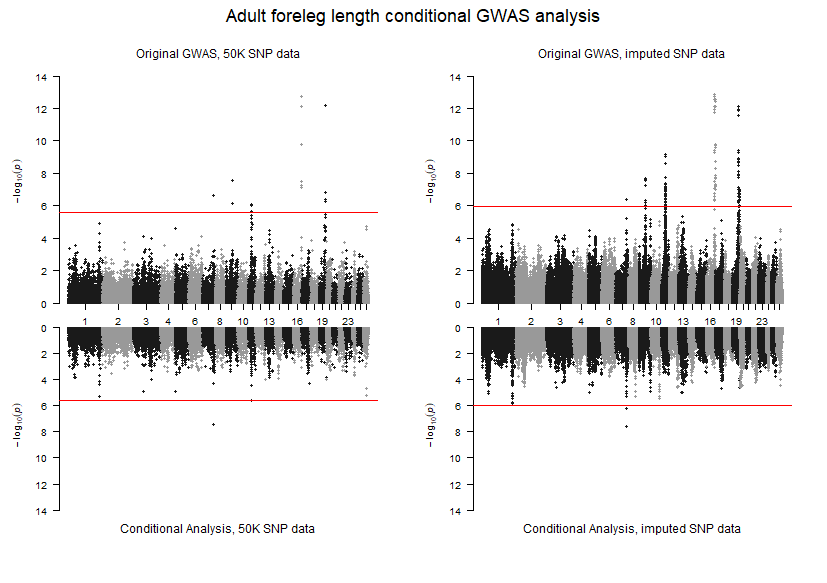

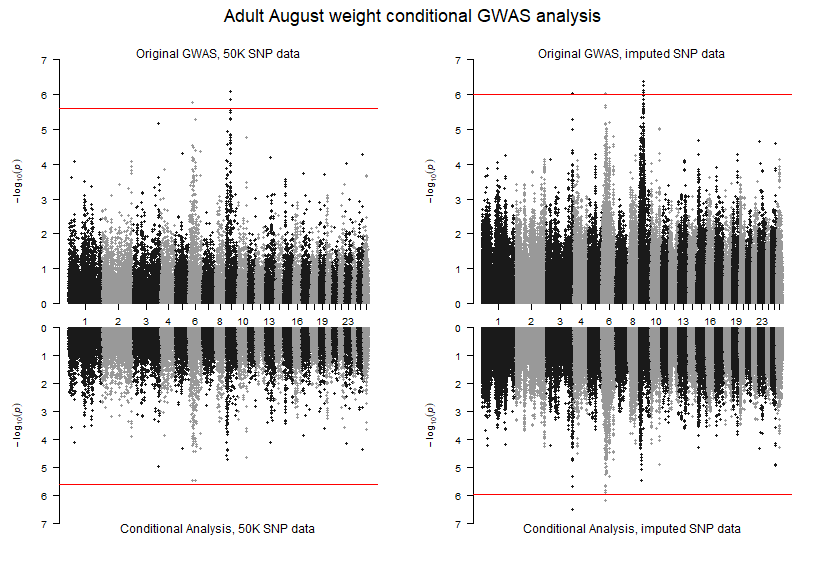

**E**

**F**

**Sup**

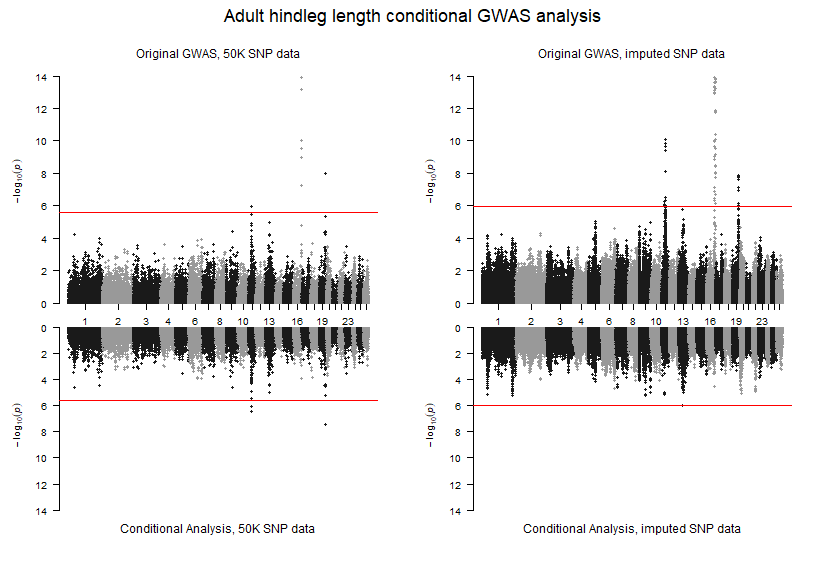

**G**

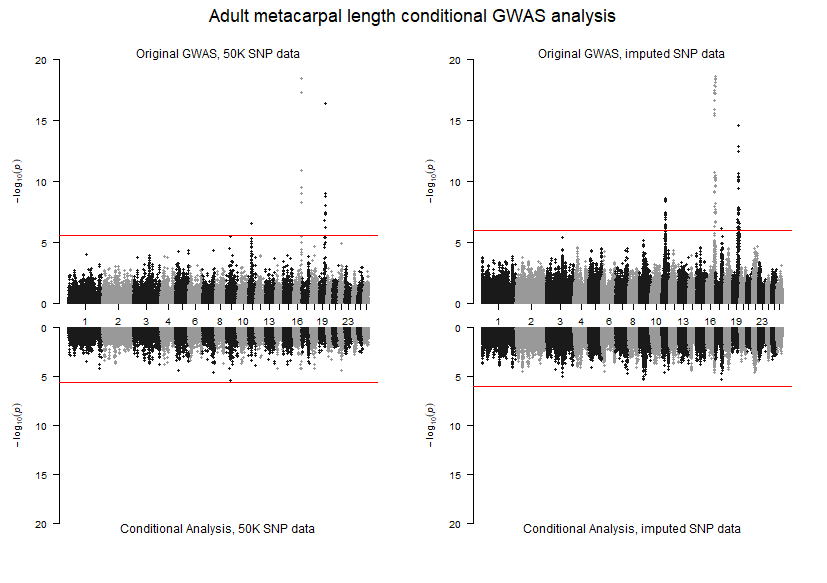

**H**

**Supplementary Figure 2** Miami plots of GWAS using the 50K SNP data (top left) and the imputed SNP data (top right) and conditional analysis using the 50K SNP data (bottom left) and the imputed SNP data (bottom right) for **A)** birth weight (imputed SNP data only), **B)** lamb foreleg length, **C)** lamb hindleg length, **D)** lamb metacarpal length, **E)** adult August weight **F)** adult foreleg length, **G)** adult hindleg length, **H)** adult metacarpal length, and **I)** adult jaw length. The red line represents the significance threshold (2.49e^-06^ for the 50K SNP data and 1.03e^-06^ for the imputed SNP data) – any SNPs above this threshold are considered to be significantly associated with variation in their respective traits.

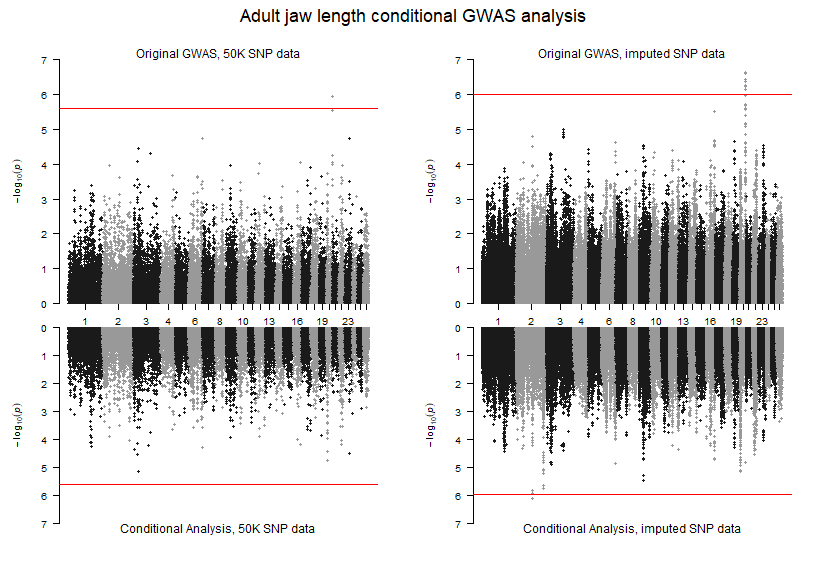

**I**
