## Supplementary Figure 3 for "The impact of SNP density on quantitative genetic analyses of body size traits in a wild population of Soay sheep"

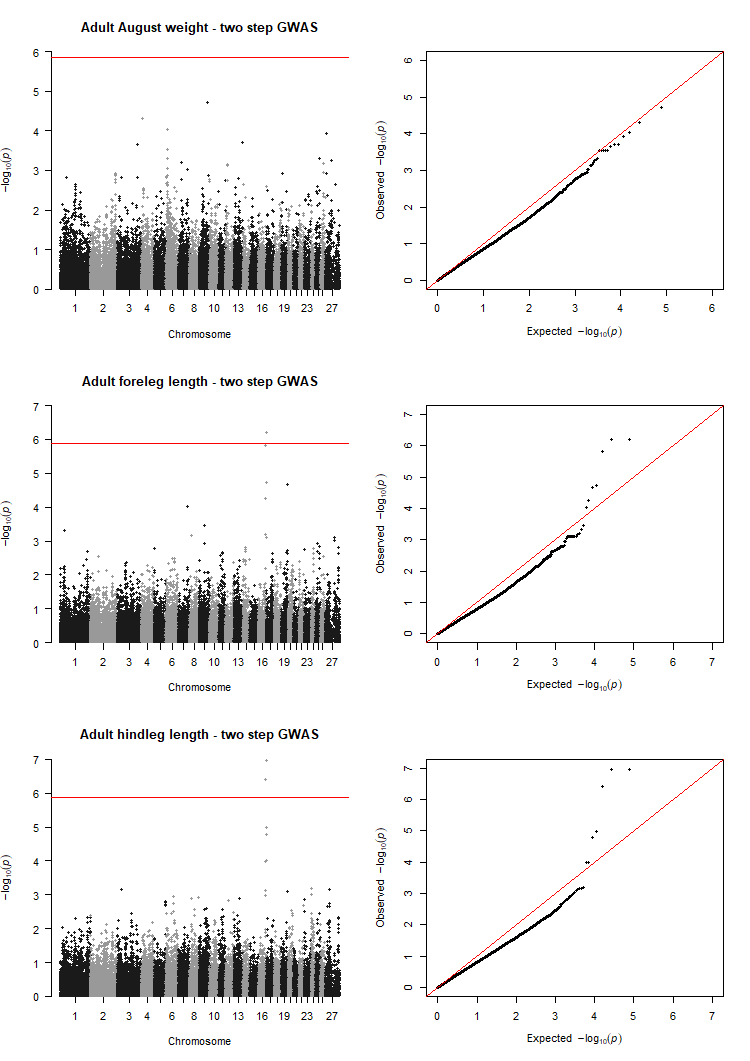

**A**

**B**

**C**

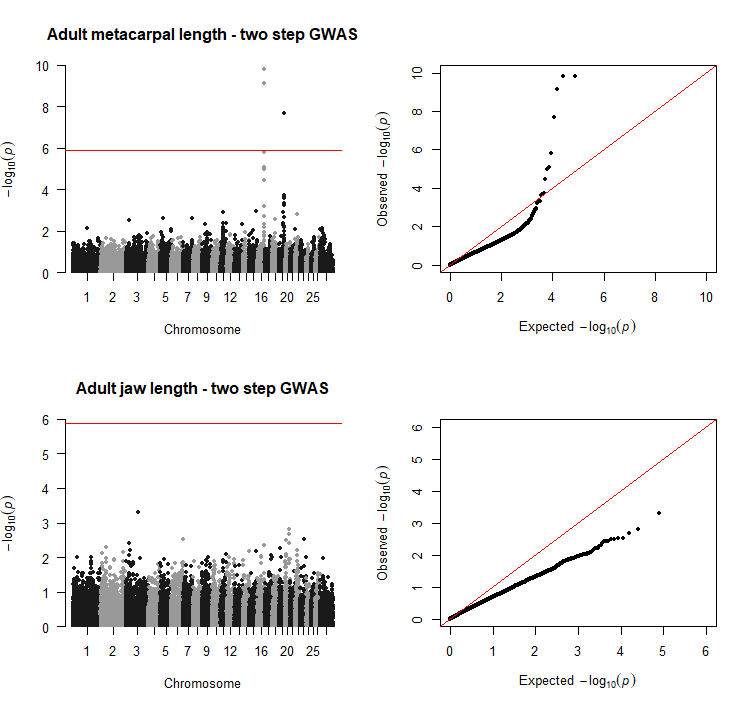

**E**

**D**

**Supplementary Figure 3** Manhattan and QQ plots using the two-step GWAS method with the 50K SNP data for the adult traits **A)** August weight, **B)** foreleg length, **C)** hindleg length, **D)** metacarpal length, and **E)** jaw length. In the Manhattan plots, the red line represents the significance threshold (1.35 × 10−6) – any SNPs above this threshold are considered to be significantly associated with variation in their respective traits. In the QQ plots, the red line is the X=Y line.
