## Supplementary Methods for "The impact of SNP density on quantitative genetic analyses of body size traits in a wild population of Soay sheep"

The program used in all analyses reported here was DISSECT. At first sight this program is not well set up to run models for repeated measures traits. Here, we describe how we overcame this issue.

In a standard GWAS, both the phenotype and the genotype files contain a Family ID column (FID) and an Individual ID column (IID) that has a unique identifier for each individual in the dataset, and links the phenotype for an individual to the correct genotype. However, in a repeatability model, an individual may have more than one record in the phenotype file, and so there will be repeated IDs in the IID column (see below).

In order to avoid having repeated IDs, a unique record identifier is used instead in the IID column, and the original IDs are included as a random effect column in order to account for permanent environment effects. In order for the phenotypes to link with the correct genotype, the IID column in the genotype file must also be replaced with the unique record identifier. As each phenotype must link to a genotype row, this means the genotype row for each individual must be duplicated to match the number of records for that individual (see below).

| **FID** | **IID** | **Phenotype** | **Year** | **REAL_ID** |
| --- | --- | --- | --- | --- |
| 1 | 12000 | 3 | 2000 | 1 |
| 1 | 12001 | 3.1 | 2001 | 1 |
| 1 | 12003 | 3.2 | 2003 | 1 |
| 1 | 22000 | 5 | 2000 | 2 |
| 1 | 22002 | 4.7 | 2002 | 2 |

| **FID** | **IID** |  |  |  |  |
| --- | --- | --- | --- | --- | --- |
| 1 | 12000 | AA | AT | GA | CC |
| 1 | 12001 | AA | AT | GA | CC |
| 1 | 12003 | AA | AT | GA | CC |
| 1 | 22000 | GA | GT | GG | AC |
| 1 | 22000 | GA | GT | GG | AC |

*The example phenotype file (left) and genotype file (right) after formatting. The IID column in both files has been changed to use a unique identifier for each record – in this case, the individual ID followed by the year. Both files also now have a row for each record for each individual.*

| **FID** | **IID** |  |  |  |  |
| --- | --- | --- | --- | --- | --- |
| 1 | 1 | AA | AT | GA | CC |
| 1 | 2 | GA | GT | GG | AC |

| **FID** | **IID** | **Phenotype** | **Year** |
| --- | --- | --- | --- |
| 1 | 1 | 3 | 2000 |
| 1 | 1 | 3.1 | 2001 |
| 1 | 1 | 3.2 | 2003 |
| 1 | 2 | 5 | 2000 |
| 1 | 2 | 4.7 | 2002 |

*An example phenotype file (left) and genotype (file) for a repeatability model before formatting. Here we have two individuals, one which has had their phenotype recorded 3 times, and one which has had their phenotype recorded twice. Year has also been included as a covariate.*
