## Supplementary Table 1 for "The impact of SNP density on quantitative genetic analyses of body size traits in a wild population of Soay sheep"

| **Age** | **Trait** | **Variance component** | **Variance - 50K SNP data** | | **Variance:Phenotypic variance ratio – 50K SNP data** | | **Variance - imputed SNP data** | | **Variance:Phenotypic variance ratio – imputed SNP data** | |
| --- | --- | --- | --- | --- | --- | --- | --- | --- | --- | --- |
| **Neonate** | **Birth weight** | GRM | 0.009 | (0.004) | 0.051 | (0.020) | 0.009 | (0.004) | 0.051 | (0.020) |
|  |  | Year of birth | 0.028 | (0.008) | 0.150 | (0.036) | 0.028 | (0.008) | 0.150 | (0.036) |
|  |  | Mother ID | 0.041 | (0.004) | 0.219 | (0.022) | 0.041 | (0.004) | 0.219 | (0.022) |
|  |  | Residual | 0.108 | (0.004) | 0.580 | N/A | 0.108 | (0.004) | 0.580 | N/A |
| **Lamb** | **Weight** | GRM | 0.493 | (0.133) | 0.091 | (0.024) | 0.472 | (0.131) | 0.087 | (0.024) |
|  |  | Year of birth | 0.912 | (0.264) | 0.168 | (0.041) | 0.910 | (0.264) | 0.168 | (0.041) |
|  |  | Mother ID | 0.828 | (0.135) | 0.152 | (0.024) | 0.825 | (0.135) | 0.152 | (0.024) |
|  |  | Residual | 3.201 | (0.147) | 0.589 | N/A | 3.214 | (0.148) | 0.593 | N/A |
|  | **Foreleg** | GRM | 8.285 | (1.467) | 0.144 | (0.027) | 8.138 | (1.448) | 0.142 | (0.027) |
|  |  | Year of birth | 16.337 | (4.590) | 0.284 | (0.058) | 16.332 | (4.590) | 0.285 | (0.058) |
|  |  | Mother ID | 4.442 | (1.029) | 0.077 | (0.019) | 4.239 | (1.024) | 0.074 | (0.019) |
|  |  | Residual | 28.480 | (1.364) | 0.495 | N/A | 28.603 | (1.370) | 0.499 | N/A |
|  | **Hindleg** | GRM | 14.080 | (2.742) | 0.152 | (0.029) | 13.536 | (2.694) | 0.146 | (0.028) |
|  |  | Year of birth | 12.262 | (3.651) | 0.132 | (0.034) | 12.333 | (3.670) | 0.133 | (0.035) |
|  |  | Mother ID | 8.967 | (2.007) | 0.096 | (0.021) | 8.792 | (2.008) | 0.095 | (0.021) |
|  |  | Residual | 57.620 | (2.642) | 0.620 | N/A | 57.966 | (2.655) | 0.626 | N/A |
|  | **Metacarpal** | GRM | 5.490 | (0.794) | 0.285 | (0.037) | 5.911 | (0.817) | 0.309 | (0.038) |
|  |  | Year of birth | 0.815 | (0.323) | 0.042 | (0.016) | 0.805 | (0.319) | 0.042 | (0.016) |
|  |  | Mother ID | 2.042 | (0.506) | 0.106 | (0.026) | 1.898 | (0.503) | 0.099 | (0.026) |
|  |  | Residual | 10.892 | (0.651) | 0.566 | N/A | 10.529 | (0.656) | 0.550 | N/A |
|  | **Jaw** | GRM | 5.033 | (0.802) | 0.243 | (0.036) | 5.272 | (0.819) | 0.256 | (0.037) |
|  |  | Year of birth | 1.065 | (0.403) | 0.052 | (0.019) | 1.057 | (0.399) | 0.051 | (0.019) |
|  |  | Mother ID | 3.210 | (0.581) | 0.155 | (0.027) | 3.161 | (0.582) | 0.153 | (0.027) |
|  |  | Residual | 11.377 | (0.667) | 0.550 | N/A | 11.122 | (0.675) | 0.540 | N/A |
| **Adult** | **Weight** | GRM | 2.582 | (0.396) | 0.251 | (0.035) | 2.539 | (0.393) | 0.247 | (0.035) |
|  |  | Year of capture | 1.311 | (0.343) | 0.127 | (0.029) | 1.299 | (0.341) | 0.127 | (0.029) |
|  |  | Permanent environment | 3.338 | (0.315) | 0.324 | (0.032) | 3.354 | (0.318) | 0.327 | (0.033) |
|  |  | Residual | 3.073 | (0.087) | 0.298 | N/A | 3.074 | (0.087) | 0.299 | N/A |
|  | **Foreleg** | GRM | 10.817 | (1.365) | 0.266 | (0.038) | 11.505 | (1.396) | 0.283 | (0.039) |
|  |  | Year of capture | 14.662 | (3.782) | 0.361 | (0.060) | 14.766 | (3.810) | 0.363 | (0.060) |
|  |  | Permanent environment | 7.806 | (0.892) | 0.192 | (0.029) | 7.105 | (0.880) | 0.175 | (0.028) |
|  |  | Residual | 7.326 | (0.219) | 0.180 | N/A | 7.327 | (0.219) | 0.180 | N/A |
|  | **Hindleg** | GRM | 19.597 | (2.444) | 0.424 | (0.042) | 20.611 | (2.487) | 0.446 | (0.042) |
|  |  | Year of capture | 1.883 | (0.631) | 0.041 | (0.013) | 1.849 | (0.622) | 0.040 | (0.013) |
|  |  | Permanent environment | 17.284 | (1.610) | 0.374 | (0.038) | 16.268 | (1.595) | 0.352 | (0.038) |
|  |  | Residual | 7.464 | (0.218) | 0.161 | N/A | 7.464 | (0.218) | 0.162 | N/A |
|  | **Metacarpal** | GRM | 10.139 | (1.141) | 0.644 | (0.047) | 10.128 | (1.130) | 0.653 | (0.048) |
|  |  | Year of birth | 0.102 | (0.110) | 0.006 | (0.007) | 0.111 | (0.111) | 0.007 | (0.007) |
|  |  | Residual | 5.496 | (0.630) | 0.349 | N/A | 5.275 | (0.634) | 0.340 | N/A |
|  | **Jaw** | GRM | 11.170 | (1.402) | 0.557 | (0.051) | 10.895 | (1.371) | 0.554 | (0.051) |
|  |  | Year of birth | 0.901 | (0.396) | 0.045 | (0.019) | 0.829 | (0.372) | 0.042 | (0.018) |
|  |  | Residual | 7.979 | (0.836) | 0.398 | N/A | 7.944 | (0.842) | 0.404 | N/A |

**Supplementary Table 1** Estimates for variance components and ratios to phenotypic variance when using the 50K SNP dataset (left) and using the imputed SNP dataset (right), with standard errors shown in brackets.
