## Supplementary Table 2 for "The impact of SNP density on quantitative genetic analyses of body size traits in a wild population of Soay sheep"

|  |  |  | **50K SNP data** | | | | | **Imputed SNP data** | | | | |
| --- | --- | --- | --- | --- | --- | --- | --- | --- | --- | --- | --- | --- |
|  |  | **Chr** | **Top SNP** | **Top SNP position** | **Top SNP MAF** | **Genetic variance explained by top SNP** | **Significant SNPs on chr** | **Top SNP** | **Top SNP position** | **Top SNP MAF** | **Genetic variance explained by top SNP** | **Significant SNPs on chr** |
| Neonate | Birth weight | 1 | N/A | N/A | N/A | 0 | 0 | **oar3_OAR1_175180452** | 175180452 | 0.1362 | 0.043615 | 4 |
|  |  | 7 | N/A | N/A | N/A | 0 | 0 | **oar3_OAR7_54963456** | 54963456 | 0.2087 | 0.059769 | 3 |
| Lamb | Foreleg | 16 | **s23172.1** | 69726554 | 0.03675 | 0.005266 | 2 | **oar3_OAR16_69873504** | 69873504 | 0.09982 | 0.011736 | 18 |
|  | Hindleg | 16 | **s23172.1** | 69726554 | 0.03701 | 0.00693 | 3 | **oar3_OAR16_69873504** | 69873504 | 0.1009 | 0.014778 | 32 |
|  | Metacarpal | 3 | OAR3_100483326.1 | 94492563 | 0.4213 | 0.020807 | 1 | OAR3_100483326.1 | 94492563 | 0.4213 | 0.020458 | 1 |
|  |  | 16 | **s22142.1** | 69679810 | 0.04144 | 0.009707 | 9 | **s22142.1** | 69679810 | 0.04144 | 0.008927 | 63 |
|  |  | 19 | **s74894.1** | 52470202 | 0.1063 | 0.023958 | 19 | **s74894.1** | 52470202 | 0.1063 | 0.022037 | 232 |
| Adult | August weight | 3 | N/A | N/A | N/A | 0 | 0 | oar3_OAR3_209859215 | 209859215 | 0.3548 | 0.029722 | 1 |
|  |  | 6 | OAR6_38919831.1 | 34800345 | 0.3544 | 0.03111 | 1 | oar3_OAR6_35627634 | 35627634 | 0.3496 | 0.030846 | 1 |
|  |  | 9 | **OAR9_37208898.1** | 35300316 | 0.4533 | 0.031326 | 2 | **oar3_OAR9_36632898** | 36632898 | 0.3397 | 0.032516 | 7 |
|  | Foreleg | 7 | s48811.1 | 84579439 | 0.1423 | 0.013069 | 1 | s48811.1 | 84579439 | 0.1423 | 0.011745 | 1 |
|  |  | 9 | **s50107.1** | 50463148 | 0.4902 | 0.02987 | 2 | **oar3_OAR9_50469115** | 50469115 | 0.4903 | 0.028574 | 13 |
|  |  | 11 | **OAR11_33427461.1** | 31380176 | 0.4232 | 0.023463 | 3 | **oar3_OAR11_30635038** | 30635038 | 0.2814 | 0.028791 | 46 |
|  |  | 16 | **s22142.1** | 69679810 | 0.04643 | 0.008024 | 7 | **s22142.1** | 69679810 | 0.04643 | 0.007635 | 61 |
|  |  | 19 | **s74894.1** | 52470202 | 0.1105 | 0.02042 | 7 | **oar3_OAR19_52340442** | 52340442 | 0.1059 | 0.018147 | 105 |
|  | Hindleg | 11 | OAR11_32693010.1 | 30811134 | 0.3913 | 0.022496 | 1 | **oar3_OAR11_30832740** | 30832740 | 0.2889 | 0.032152 | 21 |
|  |  | 16 | **s22142.1** | 69679810 | 0.04631 | 0.008834 | 7 | **s22142.1** | 69679810 | 0.04631 | 0.0084 | 60 |
|  |  | 19 | s74894.1 | 52470202 | 0.1118 | 0.01317 | 1 | **oar3_OAR19_52340442** | 52340442 | 0.107 | 0.011839 | 17 |
|  | Metacarpal | 11 | OAR11_33324109.1 | 31280002 | 0.444 | 0.011276 | 1 | **oar3_OAR11_31212644** | 31212644 | 0.4887 | 0.030098 | 27 |
|  |  | 16 | **s22142.1** | 69679810 | 0.04573 | 0.005487 | 7 | **oar3_OAR16_71129444** | 71129444 | 0.04545 | 0.01133 | 71 |
|  |  | 17 | N/A | N/A | N/A | 0 | 0 | oar3_OAR17_54802053 | 54802053 | 0.366 | 0.020203 | 1 |
|  |  | 19 | **s74894.1** | 52470202 | 0.1016 | 0.02023 | 10 | **oar3_OAR19_52459313** | 52459313 | 0.09867 | 0.018632 | 82 |
|  | Jaw | 20 | s07047.1 | 46575517 | 0.2773 | 0.020508 | 1 | **oar3_OAR20_46332617** | 46332617 | 0.243 | 0.023941 | 10 |

**Supplementary Table 2** GWAS results for both the 50K SNP data and the imputed SNP data. The table shows the top SNP (the SNP with the lowest p value) and its position for each chromosome as well as the minor allele frequency (MAF) and the proportion of genetic variance explained by the top SNP, followed by the number of significant SNPs on each chromosome. No SNP-trait associations were found for lamb August weight or for lamb jaw length. Bold indicates SNPs for each trait that were later fitted during conditional analysis.
