## Supplementary Table 3 for "The impact of SNP density on quantitative genetic analyses of body size traits in a wild population of Soay sheep"

|  |  |  | **50K SNP data** | | | **Imputed SNP data** | | |
| --- | --- | --- | --- | --- | --- | --- | --- | --- |
|  |  | **Chr** | **Top SNP** | **Top SNP position** | **Significant SNPs on chromosome** | **Top SNP** | **Top SNP position** | **Significant SNPs on chromosome** |
| Neonate | Birth weight | 2 | N/A | N/A | 0 | oar3_OAR2_81719138 | 81719138 | 9 |
| Lamb | Metacarpal | *3* | *OAR3_100483326.1* | *94492563* | *3* | *OAR3_100483326.1* | *94492563* | *16* |
| Adult | August weight | 3 | N/A | N/A | 0 | *oar3_OAR3_209859215* | *209859215* | *1* |
|  |  | 6 | N/A | N/A | 0 | *oar3_OAR6_35627634* | *35627634* | *1* |
|  | Foreleg | 7 | *s48811.1* | *84579439* | *1* | *s48811.1* | *84579439* | *3* |
|  | Hindleg | *11* | *OAR11_32693010.1* | *30811134* | *2* | N/A | N/A | 0 |
|  |  | *19* | *s74894.1* | *52470202* | *1* | N/A | N/A | 0 |
|  | Jaw | 2 | N/A | N/A | 0 | oar3_OAR2_137162126 | 137162126 | 1 |

**Supplementary Table 3** Conditional analysis results for both the 50K SNP data and the imputed SNP data. The table shows the top SNP (the SNP with the lowest p value) and its position for each associated region as well as the number of significant SNPs in each region. Italics indicate that this region was previously found to be significantly associated with the focal trait but was not fitted during the conditional analysis due to only one SNP in the region being significant.
